# 3DFAACTS-SNP: Using regulatory T cell-specific epigenomics data to uncover candidate mechanisms of Type-1 Diabetes (T1D) risk

**DOI:** 10.1101/2020.09.04.279554

**Authors:** Ning Liu, Timothy Sadlon, Ying Ying Wong, Stephen Pederson, James Breen, Simon C Barry

## Abstract

**Background:** Genome-wide association and fine-mapping studies have enabled the discovery of single nucleotide polymorphisms (SNPs) and other variants that are significantly associated with many autoimmune diseases including type 1 diabetes (T1D). However, many of the SNPs lie in non-coding regions, limiting the identification of mechanisms that contribute to autoimmune disease progression.

**Methods:** Autoimmunity results from a failure of immune tolerance, suggesting that regulatory T cells (Treg) are likely a significant point of impact for this genetic risk, as Treg are critical for immune tolerance. Focusing on T1D as a model of defective function of Treg in autoimmunity, we designed a SNPs filtering workflow called 3 Dimensional Functional Annotation of Accessible Cell Type Specific SNPs (3DFAACTS-SNP) that utilises overlapping profiles of Treg-specific epigenomic data (ATAC-seq, Hi-C and FOXP3-ChIP) to identify regulatory elements potentially driving the effect of variants associated with T1D, and the gene(s) that they control.

**Results:** Using 3DFAACTS-SNP we identified 36 SNPs with plausible Treg-specific mechanisms of action contributing to T1D from 1,228 T1D fine-mapped variants, identifying 119 novel interacting regions resulting in the identification of 51 candidate target genes. We further demonstrated the utility of the workflow by applying it to three other fine-mapped/meta-analysed SNP autoimmune datasets, identifying 17 Treg-centric candidate variants and 35 interacting genes. Finally, we demonstrate the broad utility of 3DFAACTS-SNP for functional annotation of any genetic variation using all common (>10% allele frequency) variants from the Genome Aggregation Database (gnomAD). We identified 7,900 candidate variants and 3,245 candidate target genes, generating a list of potential sites for future T1D or autoimmune research.

**Conclusions:** We demonstrate that it is possible to further prioritise variants that contribute to T1D based on regulatory function and illustrate the power of using cell type specific multi-omics datasets to determine disease mechanisms. The 3DFAACTS-SNP workflow can be customised to any cell type for which the individual datasets for functional annotation have been generated, giving broad applicability and utility.

## Background

Autoimmune diseases are chronic inflammatory disorders caused by a breakdown of immunological tolerance to self-antigens, which results in an imbalance between multiple immune cells, including conventional T cells (Tconvs) and regulatory T cells (Tregs) (1). The imbalance of immune cell function can lead to the destruction of host tissues, such as is observed in multiple autoimmune diseases, including rheumatoid arthritis (RA) (joint tissues), multiple sclerosis (MS) (myelinated nerves) and inflammatory bowel disease (IBD) (intestine/colon). In the case of Type 1 Diabetes (T1D), a reduction of Treg cell function contributes to unrestrained immune destruction of the insulin-generating pancreatic beta cells (2).

Regulatory T cell function is mediated by expression of the Foxhead Box Protein 3 (FOXP3) transcription factor (TF) as evidenced by severe autoimmune diseases observed in FOXP3-deficient scurfy mice (3) and IPEX in humans (4–6). RNA sequencing and chromatin immunoprecipitation (ChIP) studies have uncovered an extensive FOXP3-dependent molecular program involved in Treg cell development and stability (7,8), and functional fitness of Treg is dependent on stable robust expression of FOXP3, such that reduced FOXP3 expression is linked to reduced Treg function. For example, in a small T1D cohort study, we have shown that there is a decrease in FOXP3 expression in the Treg of children over the first 9 months post diagnosis (9). However, since FOXP3 itself is not mutated in autoimmune diseases other than IPEX, the loss of FOXP3 levels and functional fitness is likely caused by perturbation of the Treg gene regulatory network. Hence, by decoding the regulatory network of FOXP3, and mapping the genetic risk to the key functional genes it impacts, we will gain a better understanding of how autoimmune diseases like T1D could be countered.

T1D occurs spontaneously in approximately 80% of individuals, however predisposition to the disease has a strong pattern of inheritance (10). Genome-Wide Association Studies (GWAS) have identified over 50 loci that are strongly associated with T1D, based on the genotyping of a total of 9934 cases and 16956 controls from multiple cohorts and resources (11). In addition, fine-mapping of immune-disease associated loci represented on the Immunochip Array (12) followed by a Bayesian approach identified 44 significant T1D-associated Loci and over 1,000 credible SNPs (13). While alterations in either the effector or regulatory arms of the immune system can result in loss of tolerance and autoimmune disease, we have used a Treg centric view of loss of tolerance. This is based on the observation that defects in Treg function have been reported in autoimmune diseases including T1D and MS (14,15) and that experimental deletion of FOXP3 or reduced Treg function results in autoimmune disease in many model systems (16,17).

Although GWAS have revealed significant associations between genetic variants and T1D, the vast majority of the sampled single nucleotide polymorphisms (SNPs) are located in non-coding regions that do not alter the amino acid sequence in a protein, making it difficult to assign direct biological functions to variants (18–20). Non-coding variants can be linked to direct changes in gene expression by identifying expression quantitative trait loci (eQTL) that aim to associate allelic changes to a cis (within 1Mbp of the associated gene) and trans (>1Mbp) change in gene expression (21,22). This additional direct gene expression association however still fails to identify direct mechanisms by which a specific genetic variant can change gene expression. In addition, usage of eQTLs to establish direct changes from GWAS variants is somewhat limited to local, or cis-eQTLs (23,24), whereas mounting evidence shows that long-range regulatory connections, driven by three-dimensional chromatin interactions (25,26), can mediate these changes in expression.

With the increasing affordability and availability of high-throughput sequencing techniques and various epigenomics sequencing data protocols, the impact of genome organization and accessibility can now be added to the functional annotation of genetic risk. Chromatin immunoprecipitation sequencing (ChIP-seq) allows us to identify the binding sites of a transcription factor; assay for transposase-accessible chromatin sequencing (ATAC-seq) data offers the ability to identify highly accessible regions of the genome; and high resolution chromosome conformation capture sequencing (Hi-C) data can facilitate the investigation of the three-dimensional structure of the genome. Since it is believed that the mechanisms by which non-coding SNPs contribute to diseases are mostly via changes to the function of regulatory elements (20), we believe that combining multiple genomics and epigenetics sequencing data can further reveal the relationship between GWAS SNPs and disease pathways. Our hypothesis is that the genetic variation that specifically alters Treg function will reside in open chromatin in Treg cells that is bound by FOXP3 and the genes controlled by these by regulatory regions can be identified by chromosome conformation capture approaches. Therefore, in this paper, we describe a filtering workflow using multiple sequencing data from human Tregs, aiming to identify plausible immunomodulatory mechanisms and potentially find previously unknown connections between causative variant SNPs significantly associated with T1D and the genes they impact.

## Methods

### Cell preparation

Peripheral blood mononuclear cells (PBMCs) were isolated from whole blood obtained from healthy human donors with informed consent at the Women’s and Children’s Hospital, Adelaide (ethics approval and consent see Declarations section). Cells were labelled with the following fluorochrome conjugated anti-human monoclonal antibodies: anti-CD4 (BD Biosciences, BUV395 Mouse Anti-Human), anti-CD25 (BD Biosciences, BV421), anti-CD127 (BD Biosciences, PE-CF594) and viability dye (BD Biosciences, BD Horizon Fixable Viability Stain 700) for FACS analysis by surface expression staining. Regulatory T (Treg) cells were sorted as CD4+ CD25hi CD127dim population (>90% purity). Following cell sorting Treg cells were plated at 100,000 cells per well in a 96-well U-bottom plate and maintained in complete X-VIVO 15 culture media (X-VIVO 15 Serum-free media supplemented with 2 mM HEPES pH 7.8, 2 mM L-glutamine and 5% heat inactivated human serum) in 400U/mL rIL-2 for 2 hours at 37oC in a humidified 5% CO2 incubator prior to cell preparation for ATAC-seq experiment.

### ATAC-seq library preparation and high-throughput sequencing

Treg cells were rested for 2-hour post sort and then were either left untreated or stimulated with beads conjugated with anti-CD3 and anti-CD28 antibodies (Dynabeads Human T-Expander CD3/CD28, Gibco no. 11141D, Life Technologies) in complete X-VIVO 15 culture in 400U/mL rIL-2 at a cell/bead ratio of 1:1 for 48 hours. After 48 hours Dynabeads were removed from culture medium by magnetic separation. Omni ATAC-seq was then performed as described previously (27) with minor modifications. Briefly, cells with 5-15% dead cells were pretreated with 200U/µL DNase (Worthington) for 30 minutes at 37°C prior to ATAC-seq experiments. Treg cells (50, 000) were lysed in 50µL of cold resuspension buffer (RSB: 10 mM Tris-HCl pH 7.4, 10 mM NaCl, and 3 mM MgCl2) containing 0.1% NP40, 0.1% Tween-20, and 0.01% digitonin on ice for 3 minutes. The reaction was then washed with 1mL of ATAC-seq RSB containing 0.1% Tween-20 by centrifugation at 500 xg for 10 minutes at 4°C and the nuclei were resuspended in 50µL of transposition mix (30µL 2× TD buffer, 3.0µL Tn5 transposase, 16.5µL PBS, 0.5µL 1% digitonin and 0.5µL 10% Tween-20) (Illumina Inc). The transposition reaction was incubated at 37°C for 45 minutes in a thermomixer with 1000 rpm mixing. The reaction was purified using a Zymo DNA Clean & Concentrator-5 (D4014) kit. All libraries were amplified for a total of 9 PCR cycles and size selection was carried out to enrich for a fragment size window of 200 to 900bp prior to sequencing. Libraries were quantified by PCR using a KAPA Library Quantification Kit for NGS (KAPA Biosystems, Roche Sequencing). Barcoded libraries were pooled and sequenced on a paired-end 75-cycle Illumina NextSeq 550 High-Output platform (Illumina) to an average read depth of 37.1 million reads (± 4 million) per sample.

### Treg sample preparation, Hi-C library production and high-throughput sequencing

Cord blood was obtained with informed consent at the Women’s and the Children’s Hospital, Adelaide (HREC1596; WCHN Research Ethics Committee). Mononuclear cells were isolated from cord blood postpartum as previously described (28). Briefly, cord blood CD4^+^CD25^+^(Treg) were isolated from purified mononuclear cells using a Regulatory CD4^+^CD25^+^T Cell Kit (Dynabeads; <u>Invitrogen</u>, Carlsbad, CA). Ex vivo expansion of isolated T cell populations (1 × 10^6^ cells per well in a 24-well plate) were performed in X-Vivo 15 media supplemented with 5% human AB serum (Lonza, Walkersville, MD), 20 mM HEPES (pH 7.4), 2 mM L-glutamine, and 500 U/ml recombinant human IL-2 (<u>R&D Systems</u>, Minneapolis, MN) in the presence of CD3/CD28 T cell expander beads (Dynabeads; <u>Invitrogen</u>; catalogue no. 111-41D) at a bead-to-cell ratio of 3:1. Cell harvesting, Formaldehyde cross-linking (2%) and nuclei isolation was per (29,30). Treg cell nuclei were frozen in aliquots of 1×10^7^. The in situ Hi-C procedure was carried out as per Rao et al, (2014) (31) with the following modifications MboI digestion was carried out in CutSmart® Buffer (NEB) and biotin-14-dCTP (Invitrogen; catalogue no. 19518018) replaced biotin-14-dATP in the reaction to end-fill MboI overhangs. To generate DNA suitable for library construction ligated DNA in TE buffer (10mM Tris-HCL, pH8.0 and 0.1mM EDTA, pH 8.0) was sheared to an average size of 300-500bp using a Covaris S220 (Covaris, Woburn, MA) instrument with the following parameters; 130ul in a microTube AFA fibre, 140 peak incidence power, 10% Duty cycle 10%, 200 cycles per burst for 55 seconds. Sheared fragment ends were made suitable for adapter ligation with a NEBNext^®^ Ultra II End Repair/dA-Tailing Module (NEB #<u>E7546</u>). For adapter ligation the End Prep reaction was split into two and appropriately diluted NEBNext Adaptor ligated to fragment ends using the NEBNext Ultra II Ligation module. Hi-C libraries were split between 5 separate PCR reactions and directly amplified off the T1 beads using NEBNext Index Primers (set 1) and the NEBNext® Ultra™ II Q5® Master Mix. Library size distribution was determined using an Experion DNA 1K kit and library concentration estimated by real time qPCR using a Kapa universal Library quantitation kit (Roche Sequencing Solutions; 07960140001). Hi-C libraries were sequenced on a Illumina NextSeq 500 Mid-output platform (2x 150bp).

### ATAC-seq data analysis

The sequencing data quality was determined using *FastQC* (ver. 0.11.7) (32) followed by trimming of Nextera adapters using *cutadapt* (ver. 1.14) (33). Trimmed reads were aligned to the human hg19 genome using *Bowtie2* (ver. 2.2.9) (34) with ‘-X 2000’ setting. For each sample quality trimming was performed with option ‘-q 10’ with unmapped and non-primary mapped reads filtered with option ‘-F 2828’ using *Samtools* (ver. 1.3.1) (35). PCR duplicates were then removed from Uniquely mapped paired reads using *Picard* (ver. 2.2.4). Mitochondrial reads, reads mapping to ENCODE hg19 blacklisted regions and mitochondrial blacklisted regions were filtered out using *BEDTools* (ver. 2.25.0). For peak calling the read start sites were adjusted to represent the center of Tn5 transposase binding event. Peaks were called from ATAC-seq data using *MACS2* (ver. 2.1.2) (36) and HINT-ATAC (37) was used to call footprints from the ATAC-seq peaks with parameters ‘--atac-seq --paired-end -- organism=hg19’.

The peak summits from resting and stimulated Treg were concatenated and sorted by chromosome and then by position. The sorted peak summits were then handled using an in-house Python script *ATACseqCollapsing.py*, which adapted a peak processing approach described by Corces et al (27) to generate a list of non-redundant peaks. Briefly, through an iterative procedure, the peak summits are extended by 249 bp upstream and 250 bp downstream to a final width of 500 bp. Any adjacent peak that overlaps with the most significant peak (significance value defined by *MACS2*) within the interval is removed. This process iterates to the next peak interval resulting in a list of non-redundant significant peaks.

### Hi-C data analysis

The raw sequencing read files were first processed using *AdapterRemoval* (ver 2.2.1a) (38) with default settings. The trimmed data were then analysed using *HiC-Pro* (ver 2.9.0) (39) with hg19 set as the reference genome and the GATCGATC as a potential ligation site. The valid interaction pairs of two technical replicates, which were stored in *allValidPairs* files were then concatenated into a single file followed by sorting based on the left interaction anchoring position of each interaction pair. The sorted interaction pairs were then processed using an in-house python script *allvalidpair2collapsingint.py* to generate non-redundant interactions. Similar to the merging process of ATAC-seq peaks, this is done by an iterative process, two anchor points of the first interaction pair are extended into windows with desired window sizes (in this case is 2kb), the following interaction pair is removed only if both anchor points are within the previous interacting window, otherwise new interaction windows are generated, and the number of removed interaction pairs of each iteration are counted, resulting in non-redundant interaction pairs with window size of 2kb. The merged interaction file was then processed using the functions *build_contact_map* and *ice_norm* from *HiC-Pro* to generate a normalised *n*n* matrix for subsequent visualisations.

### Topologically-associated domain identification

The valid interaction pairs of two technical replicates were concatenated together, followed by mapped to equal-size bins (40kb) of the hg19 genome and normalised using *ICE* (39), resulting in a normalised interaction matrix. The matrix was then used as input to identify topologically-associated domains (TADs) via *TopDom* (40) with window size of 5.

### Visualisation & Downstream analyses

Gene set enrichment analysis (GSEA) was performed using function *enrichr* from the R package *clusterProfiler* (41) with the hallmark gene sets from Molecular Signatures Database (MSigDB). Gene ontology (GO) analysis was performed using the R package *clusterProfiler* (41), with 0.01 as P-value threshold and 0.05 as adjusted P-value threshold (Benjamini-Hochberg adjusted). Visualisation of normalised Hi-C interaction matrices (Figure 2, 3, and 5, Additional file1: Figure S2-13) was performed on 40kb resolution using an in-house R function *hicHeatmap*. The visualisations of individual filtered T1D-associated SNP loci (Figure 2, 3, 5 and Additional file1: Figure S2-14) were constructed using the R packages *Gviz* (42), *GenomicInteractions* (43) and *coMET* (44). Visualisation of the GSEA network was performed using the R package *ggraph* (45).

**Figure 1:**
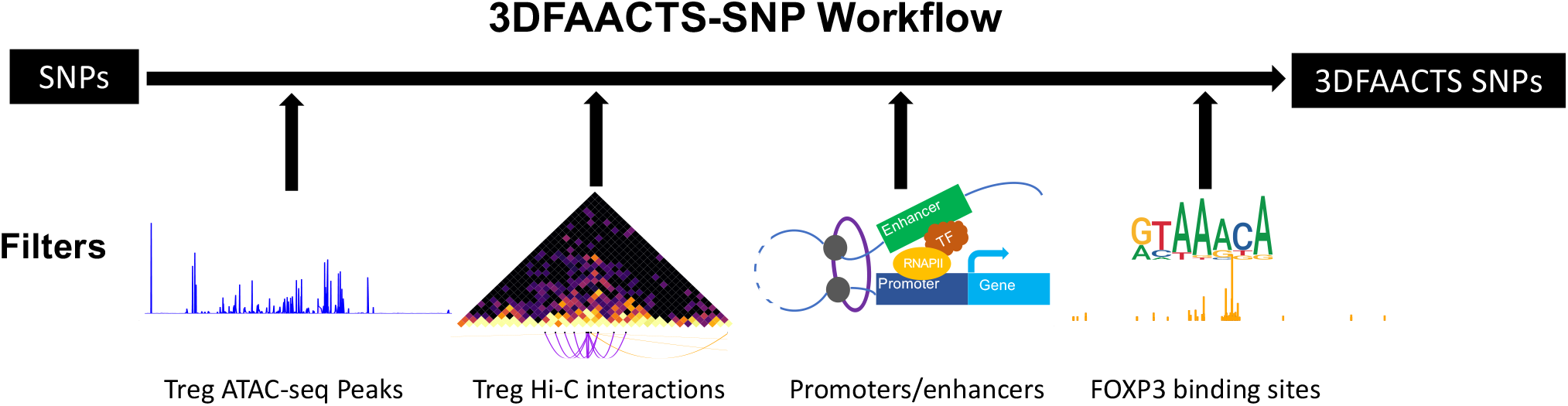
Diagram of the individual components of the Treg-specific 3DFAACTS-SNP filtering workflow for identifying variants that are potentially causative to Type 1 Diabetes (T1D). GWAS or fine-mapped variants (on the left) are intersected with different filtering elements, including Treg ATAC-seq peaks, interactions from Treg Hi-C, promoters or enhancers and previously identified FOXP3 binding regions in Treg cells (47), resulting in filtered variants we termed 3DFAACTS SNPs.

**Figure 2:**
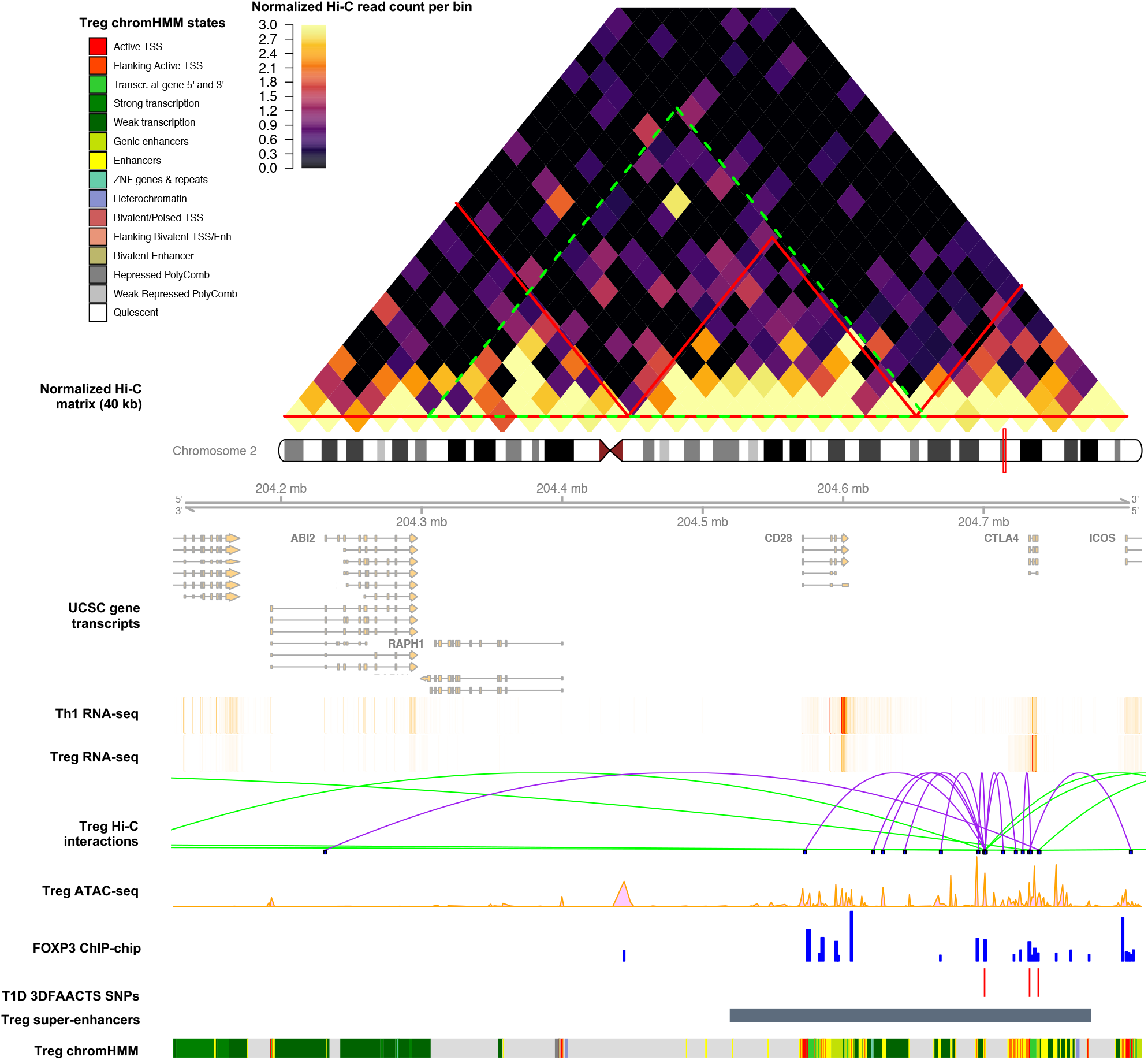
Visualisation of the CTLA4 region of filtered T1D SNPs on chromosome 2. Heatmap shows the Tregs Hi-C normalised interaction matrix (resolution of 40kb) on chr2: 203922714-205092714. The red triangles indicate Topologically Associated Domains (TADs) and the large green-dotted triangle indicates the boundary of the current plot. Tracks displayed below the chromosome 2 ideogram display workflow datasets (filtered SNPs, FOXP3-binding sites and Treg ATAC-seq and Hi-C interactions) along with various types of cell type-specific data including UCSC Gene Transcript information, T cell subsets (Thelper1 and Treg) expression data, Treg super-enhancer sets and 15-state ChromHMM track. T1D 3DFAACTS SNPs within this region are rs12990970, rs231775 and rs3087243 (from left to right).

**Figure 3:**
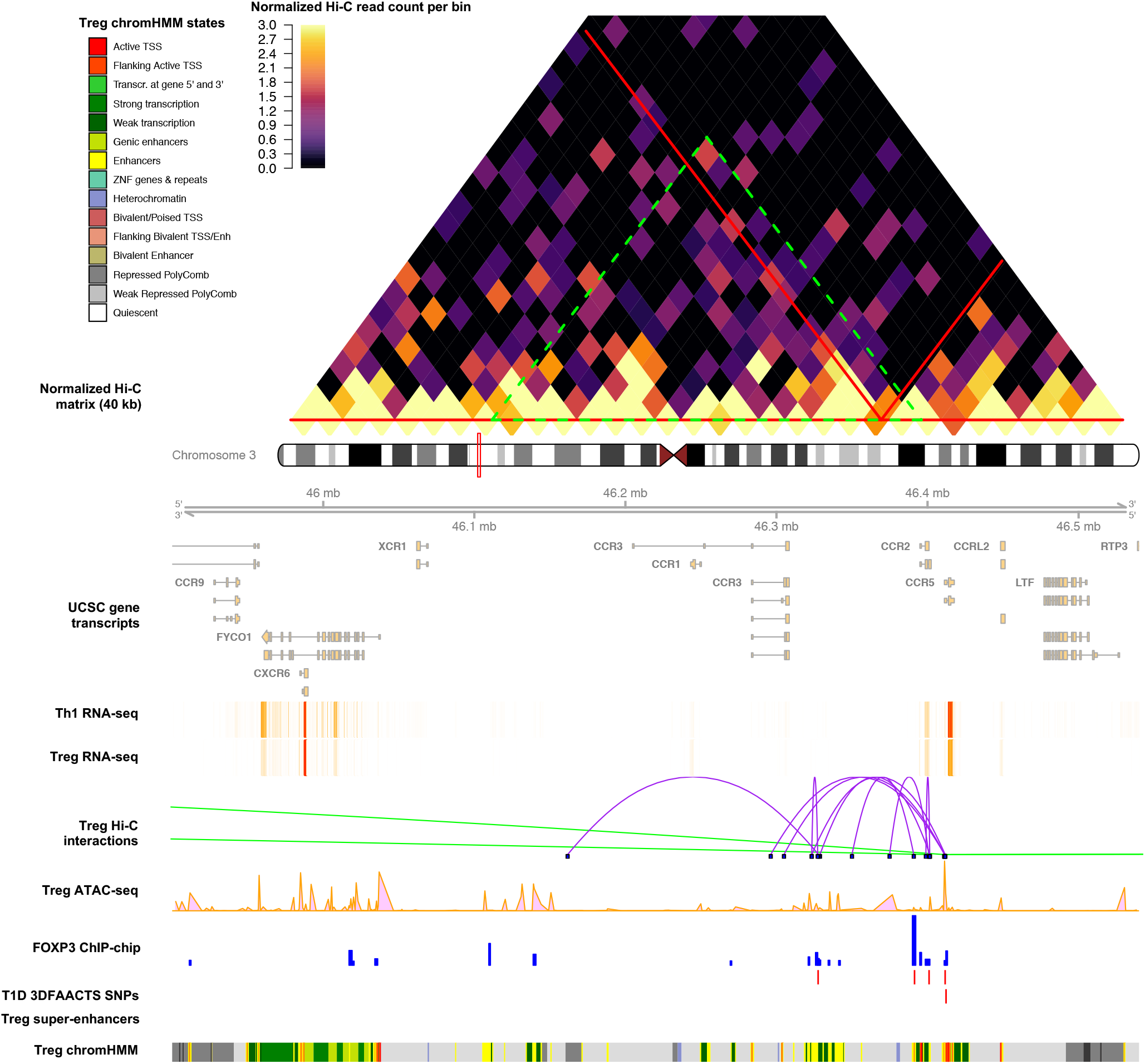
Visualisation of the CCR3/2/5 region of filtered T1D SNPs on chromosome 3. Heatmap shows the Tregs Hi-C normalised interaction matrix (resolution of 40kb) on chr3: 45600000-46840000. The red triangles indicate Topologically Associated Domains (TADs) and the large green-dotted triangle indicates the boundary of the current plot. Tracks displayed below the chromosome 3 ideogram display workflow datasets (filtered SNPs, FOXP3-binding sites and Treg ATAC-seq and Hi-C interactions) along with various types of cell type-specific data including UCSC Gene Transcript information, T cell subsets (Thelper1 and Treg) expression data, Treg super-enhancer sets and 15-state ChromHMM track. T1D 3DFAACTS SNPs within this region are rs11718385, rs6441972, rs3138042, rs2856758 and rs1799988 (from left to right).

**Figure 4:**
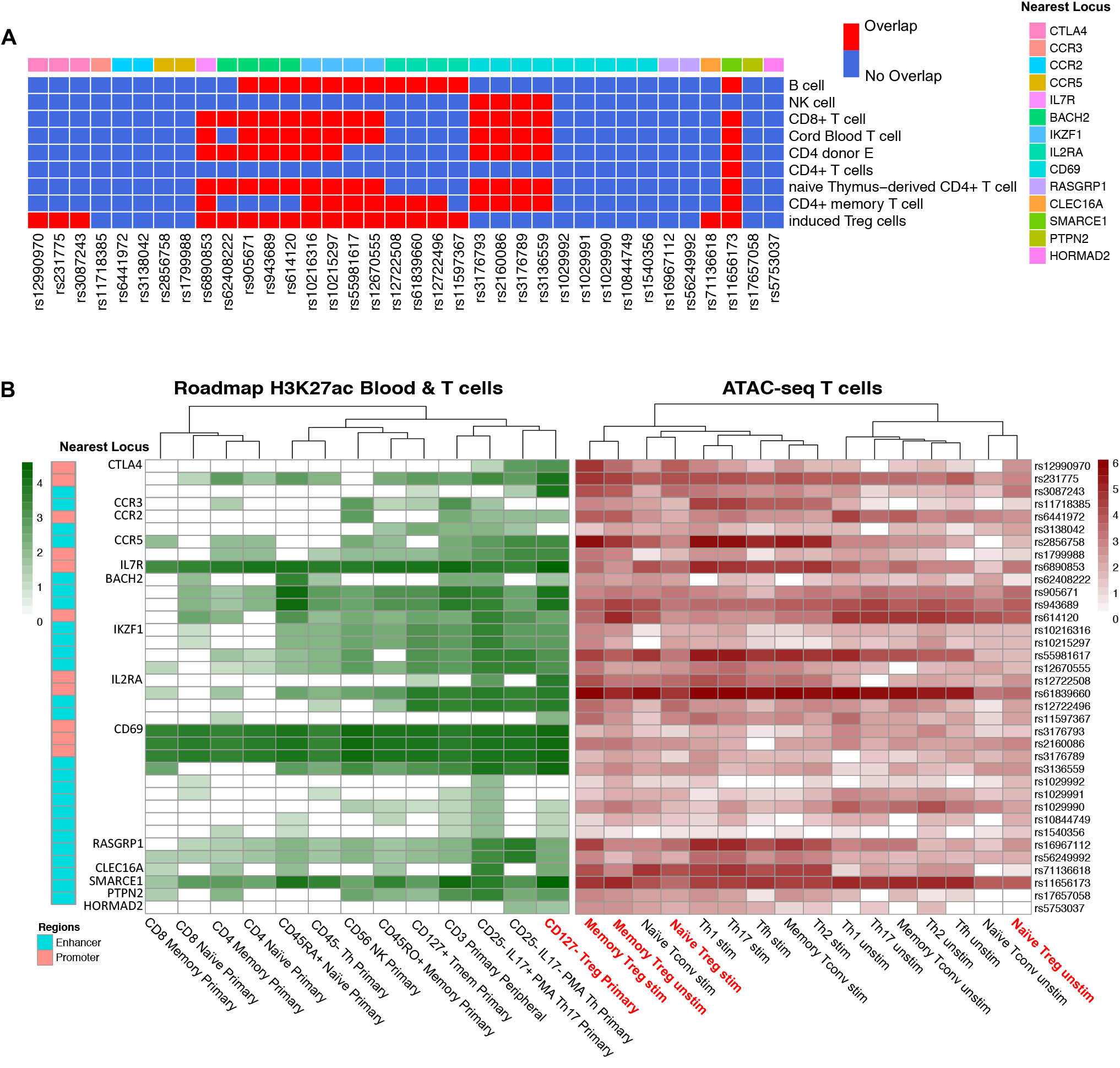
Integrating T1D 3DFAACTS SNPs with different data of T cell lineages. A. Heatmap showing overlapping status between T1D 3DFAACTS SNPs and super-enhancers of different T cell lineage from SEdb (76), where red indicates variants overlapping with SEs and blue indicates not overlapping. B. Enrichment of filtered T1D variants found within H3K27ac peaks from Epigenomics Roadmap and ATAC-seq peaks from multiple T cell lineages (52). Column names in red indicates Tregs specific datasets.

**Figure 5:**
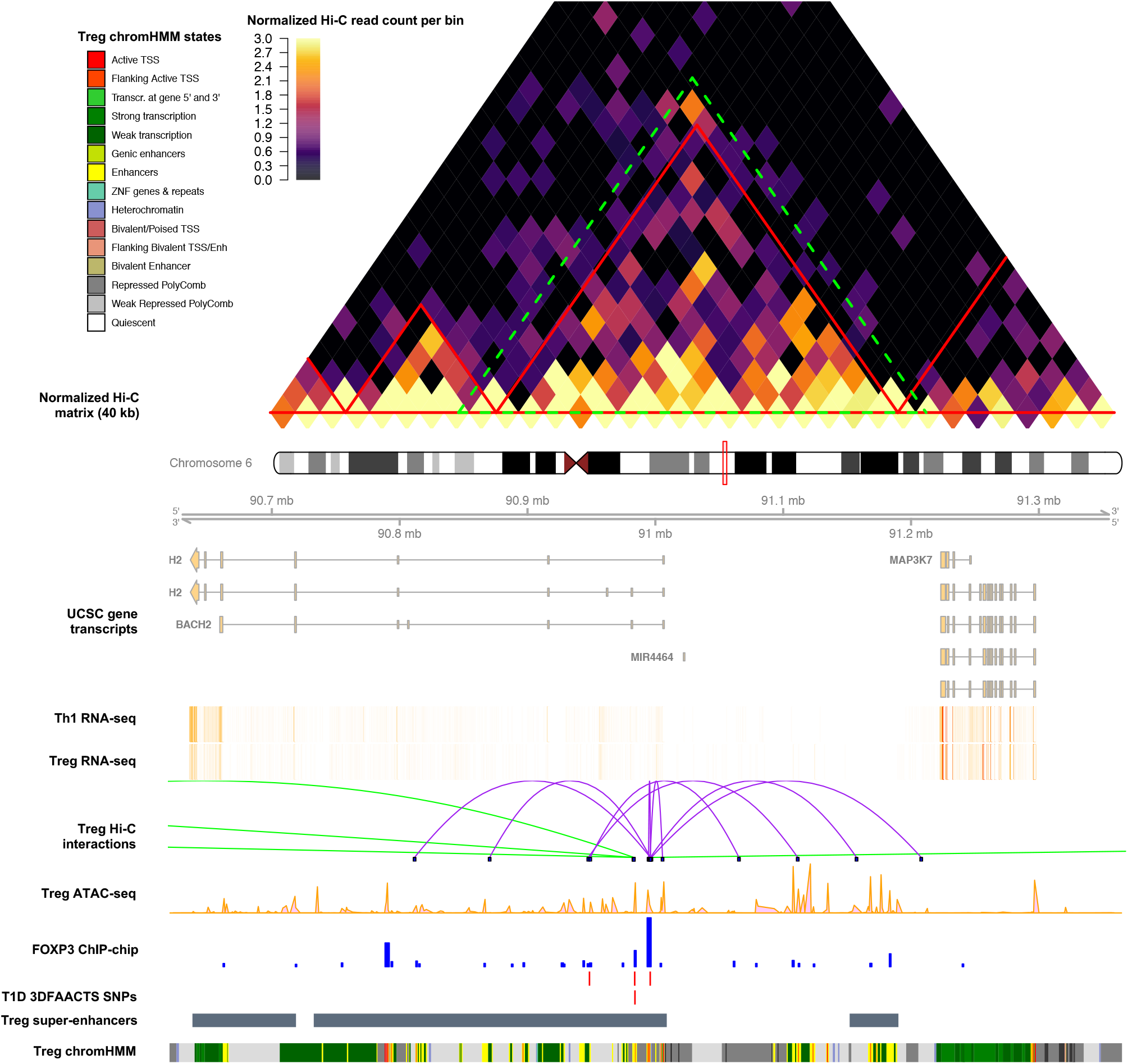
A. Visualisation of the BACH2 region of filtered T1D SNPs on chromosome 6. Heatmap shows the Tregs Hi-C normalised interaction matrix (resolution of 40kb) on chr6: 90320000-91665000. The red triangles indicate Topologically Associated Domains (TADs) and the large green-dotted triangle indicates the boundary of the current plot. Tracks displayed below the chromosome 6 ideogram display workflow datasets (filtered SNPs, FOXP3-binding sites and Treg ATAC-seq and Hi-C interactions) along with various types of cell type-specific data including UCSC Gene Transcript information, T cell subsets (Thelper1 and Treg) expression data, Treg super-enhancer sets and 15-state ChromHMM track. T1D 3DFAACTS SNPs within this region are rs62408222, rs905671, rs943689 and rs614120 (from left to right).

## Results

### Post-GWAS filtering using Treg-specific epigenomic datasets prioritises functionally relevant genetic variants contributing to T1D

As T1D is partly a consequence of Treg dysfunction, we infer that variants contained within active regulatory regions of Treg cells are likely to contribute to disease progression by impacting Treg function. A view supported by the finding that T1D associated SNPs are enriched at Treg-specific regulatory regions (46). Therefore, starting with published T1D GWAS variant information, we designed a filtering workflow (Figure 1) using multiple human Treg-specific epigenomic data to identify perturbations within defined “regulatory T cell active regions”.

In order to obtain highly accessible chromatin regions in Treg, we performed Transposase-Accessible Chromatin using sequencing (ATAC-seq) on resting and stimulated Treg cells from three donors and sequenced to an average of 37.1 million reads (± 4 million) per sample. From the ATAC-seq data, we identified 525,647 ATAC-seq peaks on average (Additional file1: Table S1). These ATAC-seq peaks were then merged into 683,954 non-redundant peaks and used to screen for variants located in accessible regions in regulatory T cells as the first filtering step of the 3DFAACTS-SNP pipeline (Figure 1).

Numerous studies have shown that three dimensional (3D) interactions play important roles in gene regulation, mediated by DNA looping bringing enhancers and promoters together at transcriptional hubs (48–50). As a result, distant loci which physically interact with disease associated regulatory regions can be potentially impacted by these regions. To identify 3D interacting regions in Treg cells, we generated and sequenced Treg *in situ* Hi-C libraries. Two technical replicates of human Treg Hi-C libraries were sequenced to an average depth of 3 million reads, and after processing using HiC-Pro (39), generated 459,244 and 1,441,362 Hi-C valid interactions respectively (Additional file1: Table S2). We extended these interactions to form 2000bp (+/- 1000bp upstream and downstream) windows at both ends of each interaction. We then collapsed interactions by merging interactions with overlapping anchors to generate non-redundant interaction pairs which represent Hi-C interactions in Tregs. These non-redundant interactions were then integrated with the variant associated ATAC-seq peaks identified above to identify accessible interacting regions.

To assign potential function to identified variant associated ATAC-seq peaks and Hi-C interacting regions we next determined the overlap of these regions with enhancer and promoter annotations. This included 113,369 enhancers (mean size of 698bp) identified by the Functional Annotation of the Mammalian Genome (FANTOM5) project (51) and promoter regions (n = 73,171) associated with GRCh37/hg19 UCSC known transcripts. Promoters were defined by extending upstream 2kb of transcription start sites (TSS). In addition, we extended the list of regulatory regions using the 15 state chromHMM model for CD4+ CD25+ CD127-Primary Treg cells from the Roadmap Epigenomics Project (52). We defined chromHMM states *EnhG*, *Enh* and *EnhBiv* as enhancers and *TssA*, *TssAFlnk*, *TssBiv* and *BivFlnk* as promoters. FANTOM5 enhancers and defined promoters and chromHMM enhancers/promoters states were then merged respectively to represent all possible genetic regulatory elements, covering 7.49 % of the genome (Additional file2: Table S3).

The transcription factor FOXP3 is critical for Treg function and orchestrating immunological tolerance, and stable high FOXP3 expression levels are observed specifically in Tregs (3,47,53). Therefore, by intersecting filtered SNPs with significant human FOXP3-binding signals, we can largely constrain SNPs within regulatory regions to FOXP3 controlled Treg-specific gene networks. We used 8,304 (mean size = 1317bp) FOXP3 ChIP-chip peaks from our previous study (47) to specify FOXP3 binding in humanTreg cells. Of interest, by searching the Gene & Autoimmune Disease Association Database (GAAD) (54), we obtained 245 annotated genes that are associated with T1D, and found a significant enrichment of FOXP3 binding sites in T1D-associated genes (Fisher exact test: P-value = 4.519e-09), suggesting a strong association between T1D risk and FOXP3 controlled Treg function. Taken together, FOXP3 binding, physical interaction, regulatory element and open chromatin regions offer a large subset of regions to use for GWAS variant prioritisation and functional annotation experiments.

### Linking fine-mapped T1D-associated variants to their targets via chromatin interactions

Genetic studies have identified over 50 candidate gene regions that contain potentially causative SNPs that impact T1D (11). Recently, a study of T1D-associated variants using Immunochip, a custom-made SNP array containing immune-related genetic variants from the 1000 genomes project (12,55), and Bayesian fine-mapping identified 1,228 putative causal variants associated with T1D (13). We used our workflow to further prioritise variants from this fine-mapped set to investigate potentially causative SNPs that contribute to T1D via affecting promoter/enhancer interaction in human Treg cells.

From the 1,228 fine-mapped T1D-associated SNPs, we identified 36 variants that meet our filtering criteria as described above, in this study we will refer to them as T1D 3DFAACTS SNPs. These variants are located at 14 different chromosomal loci and distally interact with a further 80 regions in Tregs (Table 1 & Additional file 3: Table S4). The majority of variants (71.4%, 25 out of 35 SNPs) were located in enhancer regions rather than promoters while one variant, rs614120 is located in both the *TssAFlnk* chromHMM state and T cell-specific enhancers from FANTOM5. Given that a *TssAFlnk* state can either indicate a promoter or enhancer (57), combining with the identified FANTOM enhancer information we believe that rs614120 is more likely to be located within an enhancer region.

**Table 1:**
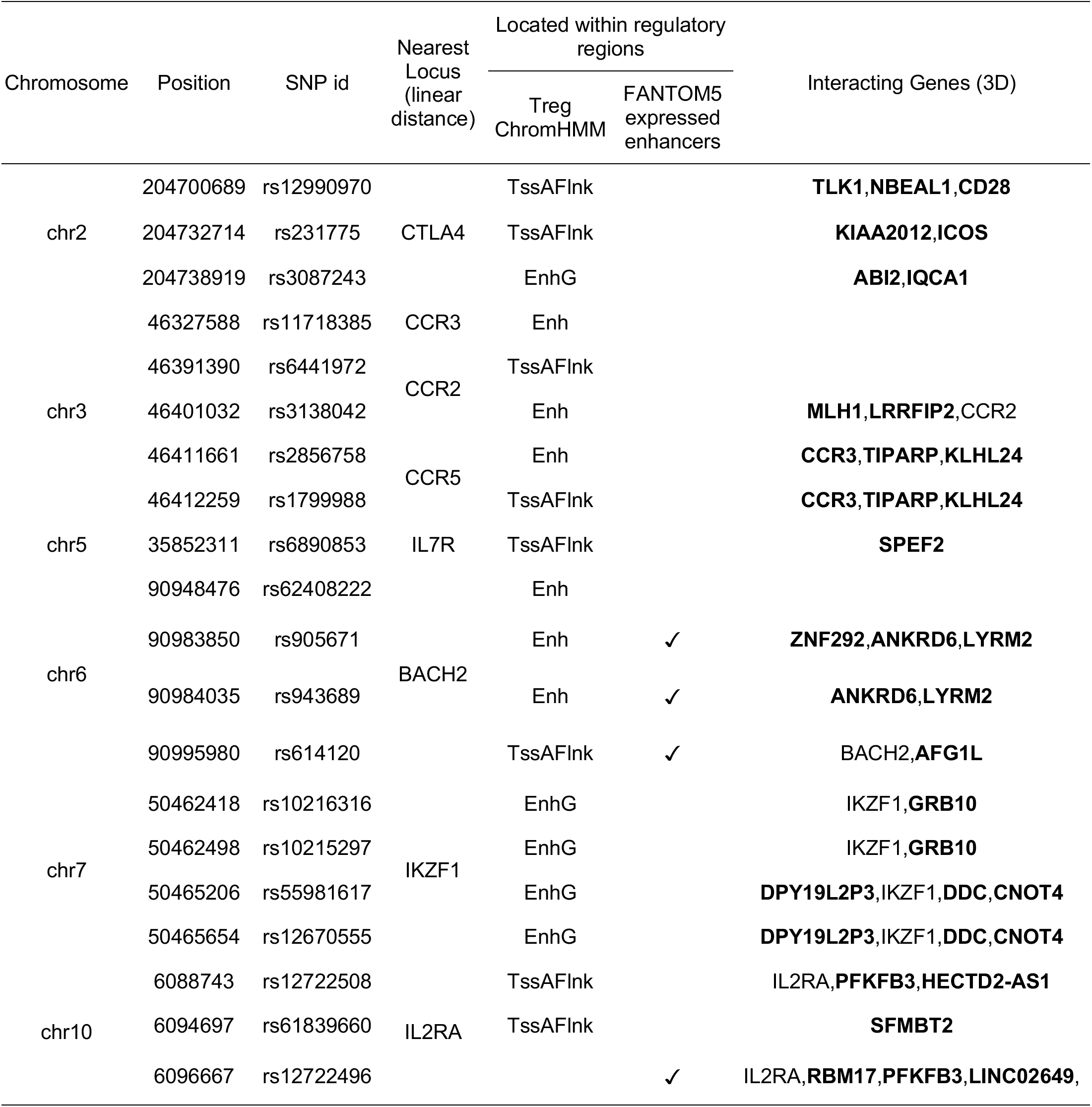
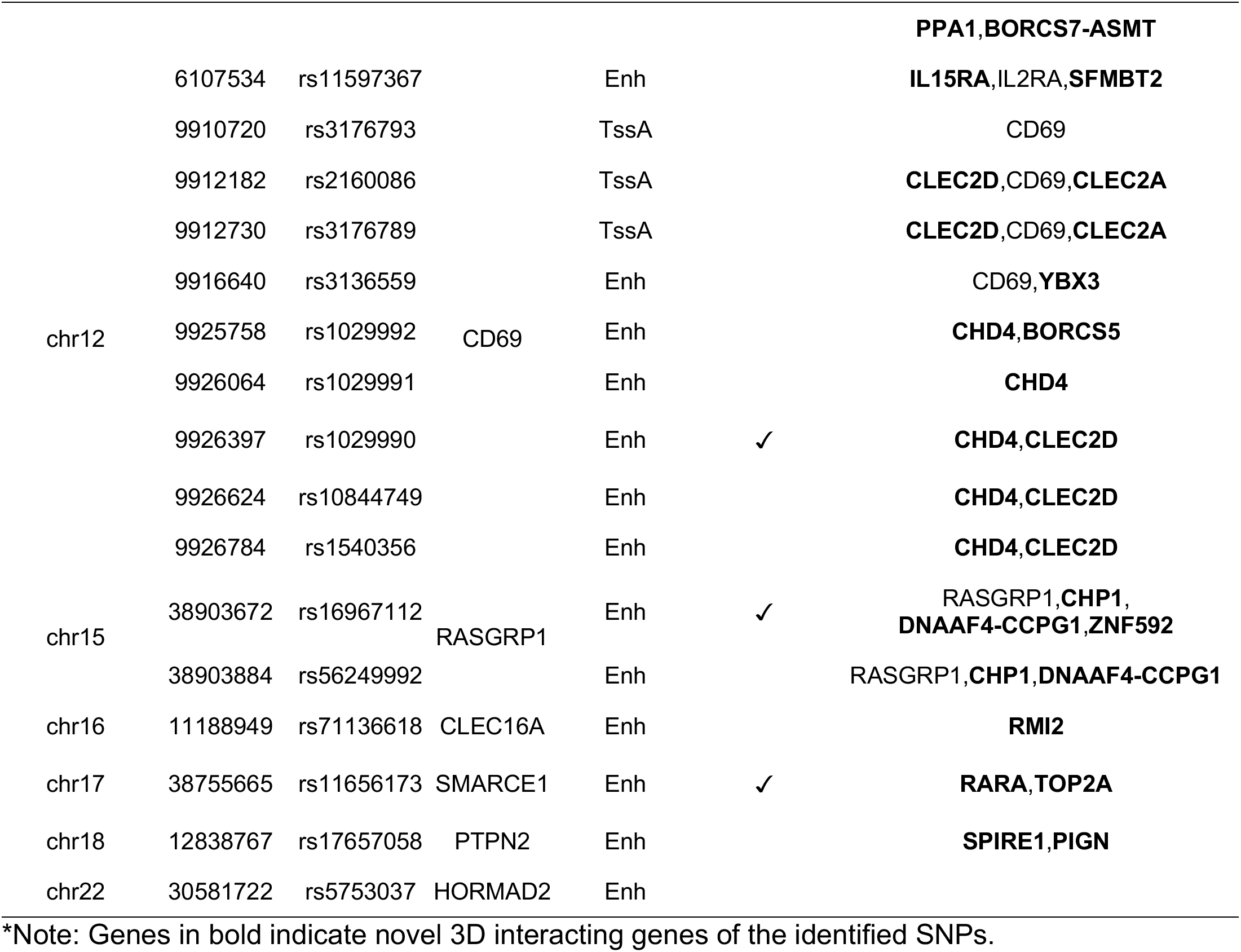
T1D 3DFAACTS SNPs identified using the 3DFAACTS-SNP filtering workflow from T1D fine-mapping SNPs (13). The nearest locus indicates the closest gene to the variants in linear distance, while 3D interacting genes are genes contact with the variants via Treg Hi-C interactions. Overlapped regulatory elements of each 3DFAACTS SNPs are displayed, including chromatin states from a 15-states model (52) and expressed enhancers from FANTOM5 (56). Detailed SNP and interaction information is contained in Supplementary Information (Additional file 3: Table S4).

Of the 14 loci identified, 8 contained more than two plausible variants across the loci. For example, variants located near the CD69 gene on chromosome 12 had the highest number of filtered variants, with 9 variants located in regulatory regions around the gene. In order to annotate the filtered variants to nearby genes, we took two approaches: annotated genes that were located in proximity to the SNPs using linear, chromosomal distances, and genes identified by their interaction with variant-containing regulatory regions via Treg Hi-C interactions (Table 1). Genes proximal to the identified 36 T1D variants include CTLA4, CCR5, IL7R, BACH2, IKZF1, IL2RA, CD69, RASGRP1, CCR3, CCR2, CLEC16A, HORMAD2 and PTPN2. These genes have previously been associated with T1D (13) and in addition other autoimmune disorders such as Multiple Sclerosis (MS), Rheumatoid Arthritis (RA), Crohn’s Disease (CD) and Inflammatory Bowel Disease (IBD) (58–62). Additionally, we annotated the filtered variants using eQTL data across all tissues from the Genotype-Tissue Expression (GTEx) project (63) and immune cells using the DICE database (64). We found that 12 filtered SNPs are annotated as the eQTL to their nearest loci (Additional file 3: Table S4) while 4 SNPs, rs11718385 (CCR3), rs62408222, rs905671 and rs943689 (BACH2) were identified as eQTL to their nearest gene in Tregs (64). These data confirmed the ability of 3DFAACTS-SNP to identify potential disease associated regulatory region-target gene networks in a cell type specific manner.

In addition to the annotation of the 36 T1D SNPs to 14 genes in closest linear proximity, 3DFAACTS-SNP identified 119 interacting regions and a further 51 genes that interact with the variant containing regulatory regions via Treg Hi-C (Table 1 & Additional file 3: Table S4). We next used the 15 states regulatory model for CD4+ CD25+ CD127- Treg primary cells from the Roadmap Epigenomics Project (52) to annotate interacting regions. These regions most frequently overlapped active chromatin states associated with transcription and gene regulation including states associated with weak transcription (*5_TxWk*) in 30% of identified regions, enhancers (*7_Enh*) in 29%, flanking active TSS (*2_TssAFlnk*) in 21% and 13% of regions located in active TSS state (*1_TssA*) (Additional file 3: Table S4). Two genes, *DPY19L2P3* and *DDC* were then dropped from further analysis as they did not overlap active states in a Treg. Additionally, searches of the GAAD (54) indicated that 45 % (22/49) of the 3D interacting genes have been previously associated with autoimmune diseases including Rheumatoid arthritis, Multiple sclerosis, Inflammatory bowel disease and T1D (Additional file 3: Table S4). Of these 22 interacting genes, 6 have been shown to be significantly associated with T1D, including BACH2, CD28, CD69, ICOS, IL2RA and RASGRP1 (Additional file 3: Table S4). Overall, by overlapping with chromHMM active states, we found 49 genes and 80 interacting regions that are active in Tregs that are in close proximity to regulatory regions carrying TD-associated variants.

Taken together, our analysis identified 31 new T1D candidate genes that may be disrupted in Treg, and a further 18 genes that have been previously associated with T1D (13). Furthermore, 61% of these interacting regions and 13 genes overlap with induced Treg super-enhancers (SEs; http://www.licpathway.net/sedb/), consistent with these regions containing important Treg functional elements. When looking at the mean normalised expression (FPKM > 1) of genes in Treg samples in Gao et al 2019 (65), 78% of interacting genes (Additional file 3: Table S4) are expressed in Tregs, all of which were enriched for T cell specific gene ontologies (Additional file 1: Figure S1). These data indicate that distal interacting regions contain regulatory regions and genes important for Treg function and are consistent with a model in which the variant containing regulatory regions may contribute to T1D by disrupting the regulation of these distal interacting genes.

### The topological neighbourhood surrounding filtered T1D variants

We next investigated the topological neighbourhood, i.e. the presence of topologically-associated or frequently interacting domains, in which regulatory regions harbouring the filtered T1D variants reside. By establishing putative boundaries of each 3D structural domain, we are then able to characterise the coordination of contacts within a loci and how they act to control gene expression. We called topologically-associated domains (TADs) using Treg Hi-C data (Additional file 4: Table S5) used in the workflow described above and integrated with publicly available super-enhancer, chromHMM data of T cell lineages and Treg expression data (66). All data was overlapped across each locus and displayed in supplementary figures 2-13.

TADs are called based on the frequency of interactions within a region (67), with physical interactions between two loci generally decaying with increasing linear distance on the chromosome (68). Genes in the closest proximity to our filtered T1D variants (Table 1), were unsurprisingly found within the same TAD. Interestingly however, we found that interacting regions and genes identified by Hi-C were only co-located within the same TAD in ∼56.5% of cases (i.e. intra-TAD interactions), with 42.6% of interactions occurring between different TADs (inter-TAD; Additional file 4: Table S5). Indeed, the linear distance between filtered variants and their 3D interacting genes (∼12.5Mb) were on average ∼2.3Mb further away compared to the average distance of intrachromosomal interactions found in the entire Treg Hi-C dataset (∼10.2Mb), indicating that Treg-active, FOXP3-bound regions impact genes across much greater linear distances than regular connections.

A high degree of chromatin interactions between genes and enhancer regions was detected within the filtered variant containing TADs, with these interactions both confirming previously identified SNP-target combinations and indicating potential new targets for investigation. For example, 3DFAACTS-SNP identified rs12990970 (chr2:204,700,689) as a potential causative T1D SNP. In Treg cells, rs12990970 is found in a flanking active TSS (TssAFlnk) state and it is located within a Treg super-enhancer (Figure 2 & Additional file 1: Figure S2). This variant is located in a non-coding region between gene CTLA4 and CD28 and in past studies, and it has been associated with CTLA4 as it is an eQTL for CTLA4 expression in testis although not in T lymphocytes or whole blood (Additional file 3: Table S4) (11,13,63,64). Hi-C interaction signals however do not indicate that the rs12990970-containing region interacts with the CTLA4 promoter in Treg, instead Hi-C interactions indicates that this region form interactions with promoter and enhancer regions connected to the costimulatory receptor CD28 gene (Table 1 & Figure 2), a family member known to play a critical role in Treg homeostasis and function (69) suggesting CD28 is a novel target for this variant in Treg.

Another example is on chromosome 3, where Hi-C interactions indicated that the chemokine receptor genes CCR1, CCR2, CCR3, CCR5 and CCR9 (Figure 3) are extensively linked in one TAD containing all of the filtered variants, indicating that these genes may be coordinately regulated. This is supported by previous RNA Pol-II ChIA-PET work (70) that detected interactions between chemokine gene clusters during immune responses including an increase in interactions amongst the CCR1, CCR2, CCR3, CCR5 and CCR9 genes during TNF stimulation of primary human endothelial cells (70) (Additional file 1: Figure S14). Recently, CCR2, CCR3 and CCR5 have been shown to have additional chemotaxis-independent effects on Treg cells with individual studies, reporting positive roles for individual chemokine receptors on CD25, STAT5, and FOXP3 expression and Treg potency (71–73), highlighting the importance of multiple genes at this locus on Treg function.

### Filtered T1D variants are enriched at lineage specific T cell super-enhancers

SEs usually consist of a cluster of closely spaced enhancers that are defined by their exceptionally high level of transcription co-factor binding and enhancer-associated histone modifications (i.e. H3K27ac) compared to all other active enhancers within a specific cell type (74). SEs are also linked to the control of important processes such as cell lineage commitment, development and function (75). Analysing T cell SE information annotated in the Super-Enhancer Database (76) (SEdb; http://www.licpathway.net/sedb/), 8 out of the 14 variant-containing loci were found to contain filtered T1D variants located in SEs formed in various T cell lineages including Treg cells consistent with the enrichment of autoimmune-disease associated variants within T cell super enhancers reported previously (75) (Figure 4A). The loci containing the CTLA4 and CLEC16A genes were the only loci that overlapped with Treg-specific SEs. The existence of a Treg SE is consistent with the different regulation of CTLA4 in Treg cells compared with other T cell lineages (77) and a recent report linking T1D risk variants to altered CLEC16A expression in Treg (65). Five other SNPs are located within SEs in multiple T cell types including induced Treg (iTreg) suggesting the gene controlled by these SE play a broad role in T cell function. While no Treg SEs are detectable at the CD69 locus the T1D associated variants in this region overlapped with SEs formed in other T subsets. No T cell associated SEs are found in the loci containing the CCR1/2/3/5, PTPN2, RASGRP1 and HORMAD2 genes (Figure 4A).

We then investigated the level of active enhancer marks (normalised H3K27ac-binding) and chromatin accessibility (normalised ATAC-seq peak coverage) overlapping each variant from Table 1 (Figure 4B). A range of tissue restriction patterns of chromatin states were observed using the NIH Epigenomics Roadmap data with enhancers displaying in general a more cell type-restricted pattern of H3K27ac signal compared to promoters. No variant was found to be located in a regulatory region that was exclusively active in Treg cells although rs12990970, rs231775 (CTLA4), rs11597367, rs12722508 (IL2RA) and rs5753037 (HORMAD2) are associated with a restricted H3K27ac pattern that included Treg. The absence of Treg-specific enhancers is consistent with FOXP3 binding data where FOXP3 binds many enhancer regions active in other T cell lineages to modify their activity in Treg cells (78). In particular, evidence suggests FOXP3 cooperates with other Thelper-lineage specifying transcription factors to diversify Treg cells into subsets that mirror the different Th-lineages (79–81). The majority of regions associated with the variants show an increase in chromatin accessibility upon stimulation in Treg and Thelper subsets consistent with increased enhancer activity upon T cell activation however in a few instances variants are located in regions that decrease in accessibility in stimulated Treg and Thelper subsets compared with their matched unstimulated counterpart. Notably these include the variants rs905671, rs943689 and rs614120 associated with BACH2. This is consistent with the reduction in BACH2 expression in CD4 T cells as they mature, and alteration to this repression is linked to proinflammatory effector function (82). Together these data are consistent with a model in which causal variants alter the output of enhancers that respond to environmental cues (83).

### Filtered variants disrupt Transcription Factor Binding Sites (TFBS) including a FOXP3-like binding site

Fundamental to understanding the function of specific disease associated variants is the identification of the potential impact of these non-coding variants on transcription factor binding. Analysis of ATAC-seq datasets with HINT-ATAC (37), identified over 5 million active TF footprints in chromatin accessibility profiles from stimulated and resting Treg populations (Additional file 5: Table S6). By imposing the additional FOXP3 binding annotation to the footprint dataset, we identified 7 T1D-associated variants that have the potential to alter the binding of 9 TFs, suggesting the molecular mechanisms by which these variants could impact Treg function (Additional file 6: Table S7). Of these 7 SNPs, one SNP rs3176789 is located in an active TSS chromHMM state region, while the others are located either in enhancers or flanking active TSS that are associated with active enhancers, suggesting these variants might interrupt the binding of TFs to affect enhancer functions, with the potential for a network effect on multiple genes.

We then used GWAS4D (84), which computes log-odds of probabilities of the reference and alternative alleles of a variant for each selected TF motif to calculate binding affinity, to predict the regulatory effect of each variant (Supplemental Table 8). Several of the variants are predicted to alter the binding of transcription factors with known roles in Treg and other T cell lineages including Nuclear activator of T cells (NFATC2 & NFATC3, rs1029991) (85), interferon regulatory transcription factor (IRF, rs3176789) (86), myocyte enhancer factor 2 (MEF2, rs6441972 and rs3176789) and FOX (Forkhead box, rs614120) family members. In addition, variant (rs1029991) has the potential to alter the binding of YY1 recently identified as an essential looping factor involved in promoter-enhancer interactions (87). Other variants (rs1136618 and rs3176789) potentially alter the binding of the zinc finger protein ZNF384. Although expressed in T cells, the importance of ZNF384 in T cell biology has not yet been explored.

Of note, rs614120 is predicted to decrease the binding affinity of FOXA2 in this enhancer region (Additional file 6: Table S7). As FOXA2 is not expressed in the immune compartment, this SNP may interfere with the binding of another member of the forkhead class of DNA-binding proteins eg FOXP3, which is localised to this region based on our FOXP3 ChIP (Figure 5). This suggests that a model in which rs614120 impacts the expression level of BACH2 and/or AFG1L by altered binding of a FOX protein to this enhancer.

Filtered variant rs1029991, is predicted to alter the binding of NFAT family members and/or YY1 to the enhancer region which has been linked to the cell surface expressed gene CD69. In addition, our analysis indicates that this enhancer also contacts the chromodomain helicase DNA-binding domain family member 4 (CHD4), indicating that CHD4 expression may be affected by this variant. Although CHD4 (Mi-2β) has not been previously linked to autoimmune diseases by GWAS studies, it has been shown to interacts with the T1D-associated genes IKZF1 and GATA3 and to play an important role in T cell development in the thymus and in T cell polarisation in the periphery including regulatory T cell subsets (88–90), consistent with altered regulation of CHD4 having the potential to contribute to T1D. Filtered variant rs3176789 is predicted to alter IRF and/or MEF2 binding linking these transcription factors to the regulation of the CD69, and CLEC family members CLECL1 and CLEC2D. The CD69 and CLEC2D genes have previously been associated with T1D by GWAS while CLECL1 has not. However, CLECL1 is a known target gene for eQTL rs3176789 (Additional file 3: Table S4), connecting this SNP and its associated regulatory region to CLECL1 expression rather than CD69 and suggesting a possible role for disrupted CLECL1 expression in Treg in T1D. Filtered variant rs6441972 is also predicted to influence the binding of MEF2 to a regulatory region in proximity to the promoter of CCR2. This region did not appear to interact with any other distal regulatory region or gene. Consistent with this variant disrupting CCR2 expression, CCR2 is a target gene for eQTL rs6441972, indicating that rs6441972 may result in altered CCR2 expression in a Treg in T1D by interfering with MEF2 binding.

### Filtered Treg variants identified in other autoimmune diseases

The primary rationale of our filtering workflow is that autoimmune diseases like T1D are mediated by altered Treg functions. Hence, using GWAS data for other autoimmune diseases, we aimed to discover variants which potentially act by disrupting 3D gene regulation in Tregs. Similar to filtering fine-mapped T1D-associated SNPs, here we used the 3DFAACTS-SNP filtering workflow to process variants identified by Immunochip fine-mapping experiments and meta-analysis from three studies for a broad range of autoimmune and inflammatory diseases. SNPs associated with 10 autoimmune diseases were identified, representing 221 fine-mapped SNPs associated with multiple sclerosis (MS) (91); 69 SNPs identified by the meta-analysis of celiac disease (CeD), rheumatoid arthritis (RA), systemic sclerosis (SSc), and T1D(92) (which we refer to the 4AI dataset); and 244 SNPs identified by the meta-analysis of GWAS datasets for ankylosing spondylitis (AS), Crohn’s disease (CD), psoriasis (PS), primary sclerosing cholangitis (PSC) and ulcerative colitis (UC)(93) (which we refer as 5ID dataset). Applying the 3DFAACTS-SNP pipeline we identified 9, 3 and 6 filtered variants from the MS, 4AI and 5ID datasets respectively (Additional file 7: Table S8). We identified putative target genes for these disease associated variants by Hi-C interactions resulting in 24, 8 and 8 genes linked to MS, 4AI and 5ID respectively (Additional file 7: Table S8). Many of these genes have either known roles in Treg differentiation, stability and function (GATA3 and CD84, ITCH, ILIRL2 and ILST) (94–102), or altered expression in human Treg in autoimmune-disease (ICA1, SESN3 and DLEU1) (103–105) and animal models of autoimmunity (SEPTIN7 and WWOX) (106).

Of the variants identified by 3DFAACTS-SNP, one variant (rs60600003) located at a locus on chromosome 7 was found to be associated with several diseases, including MS(91), celiac and systemic sclerosis(92), suggesting at least some of its interacting genes (ICA1, HERPUD2, SEPTIN7, ELMO1, DOCK4) may contribute to a common Treg defect in these diseases (Additional file 1: Figure S15 & Additional file 7: Table S8). When compared with the 36 variants identified from our T1D dataset analysis two variants, rs61839660 on chromosome 10 and rs3087243 on chromosome 2 were also prioritised by 3DFAACTS-SNP analysis of the 5ID and 4AI datasets respectively implicating their interacting genes SFMBT2 (rs61839660), ABI2 and IQCA1 (rs3087243) in the development of these diseases. While different variants were identified in the analysis of the various disease datasets, the regulatory elements in which these variants reside can be linked by Hi-C data to common candidate target gene such as PFKFB3 (rs12722496 and rs12722508 - T1D and rs947474 - 4AI). This is consistent with the view that common mechanistic pathways underlie some autoimmune diseases, although the specific risk allele within a locus can be disease-specific (107).

Similar to the filtered T1D SNPs, the GWAS filtered variants were more likely to be located within enhancer regions rather than promoters (Table 1 & Additional file 7: Table S8), surprising given that our defined enhancers cover less of the genome than promoters (enhancers: cover 2.23% of the human genome, while promoters cover 5.27%). This is also consistent with previous studies which have demonstrated an enrichment of disease associated variants at enhancer and super enhancer regions (75,108–110). We further annotated the filtered variants from these three datasets with GTEx eQTLs and Tregs eQTLs, identifying 4 SNPs that form an eQTL with a candidate gene target identified by Hi-C interactions (Additional file 7: Table S8). This included rs7731625-IL6ST and rs60600003-ELMO1, two SNP-gene contacts and eQTL pairings identified by 3DFAACTS-SNP as potential causative Treg defects in MS (rs7731625-IL6ST) and MS, T1D, celiac and systemic sclerosis (rs60600003-ELMO1), respectively. Of particular interest is the rs7731625-IL6ST pairing as IL6ST is a common signalling receptor of the IL6 family of cytokines known to have differing effects on Treg numbers and differentiation potential (100–102). Furthermore, the IL6-LIF axis has been proposed to regulate the balance of Th17/Treg cells with changes in Il6/LIF levels proposed to play a role in MS (99) highlighting a potential molecular mechanism for how the SNP variant rs7731625 may impinge on Treg function in MS.

### Identifying new variants that are candidates for impacting autoimmune disease

Most variants identified by GWAS have small effect sizes that together only represent a fraction of the heritability predicted by phenotype correlations between relatives (111). To account for this missing heritability, various models have been proposed including a highly polygenic architecture with small effect sizes of the causal variants (112,113), rare variants with large effect size (114,115) and epistatic mechanisms including gene-gene and gene-environment interactions (116,117). As a consequence many causal variants with small effect sizes are unlikely to reach genome wide significance in current GWAS whereas rare variants are often under-represented on SNP arrays (118). Lastly the preponderance of studies utilize populations of European descent which can result in a bias for SNPs with a higher minor allele frequencies in Europeans compared to other populations potentially limiting the relevance of these SNPs to the associated traits in non-Europeans (119). As an alternative approach to identify novel putative autoimmune disease-associated SNPs independently of association studies, we sampled 1,004,570 common variants (MAF > 0.1) from the Genome Aggregation Database (gnomAD) (version 3.0) (120) as inputs to our filtering workflow. Of these 808,857 overlapped with Tregs-specific Hi-C interactions, with 135,114 of these variants were located in promoter/enhancer regions and finally, 7,900 variants were located in FOXP3 binding regions (Additional file 8: Table S9). As a demonstration how this approach may complement current GWAS, 4,379 (55.7%) of the common variants we identified in gnomAD were not included in the largest GWAS T1D dataset to date (11) (Additional file 8: Table S9).

In order to further characterise the filtered gnomAD SNPs, we used *GIGGLE* (121) to compare the regions in which filtered SNPs reside against 15 predicted chromHMM genomic states across 127 cell types and tissues from Epigenomic Roadmap (52) (Figure 6 and Additional file 1: Figure S16), identifying positive and negative enrichment scores according to overlapping sets. Interestingly, although there was strong positive enrichment signal in active Tss (*TssA*), flanking active Tss (*TssAFlnk*) and enhancers (*Enh*) states in thymus, HSC, B- and T-cell groups, an enrichment was also observed across all cell types suggesting many of the enhancer and promoter regions and by extension their target genes are broadly expressed (Additional file 1: Figure S16). Moreover, unlike the 3DFAACTS-SNP analysis of GWAS derived data where filtered SNPs were enriched in enhancer regions, gnomAD derived SNPs are approximately evenly split between enhancers and promoter regions (Additional file 8: Table S9). Similarly, low/negative enrichment of the heterochromatin (Het) state was observed in all cell types whereas other inactive states such as repressed Polycomb (*ReprPC*, *ReprPCWk*) and the quiescent (*Quies*) states exhibited a negative enrichment in lymphoid cells. Interestingly, gnomAD SNPs demonstrated a strong negative enrichment in Treg cells for the chromatin states associated with strong transcription (*Tx*) and weak transcription (*TxWk*) potentially reflecting FOXP3 transcriptional repressor function (122).

**Figure 6:**
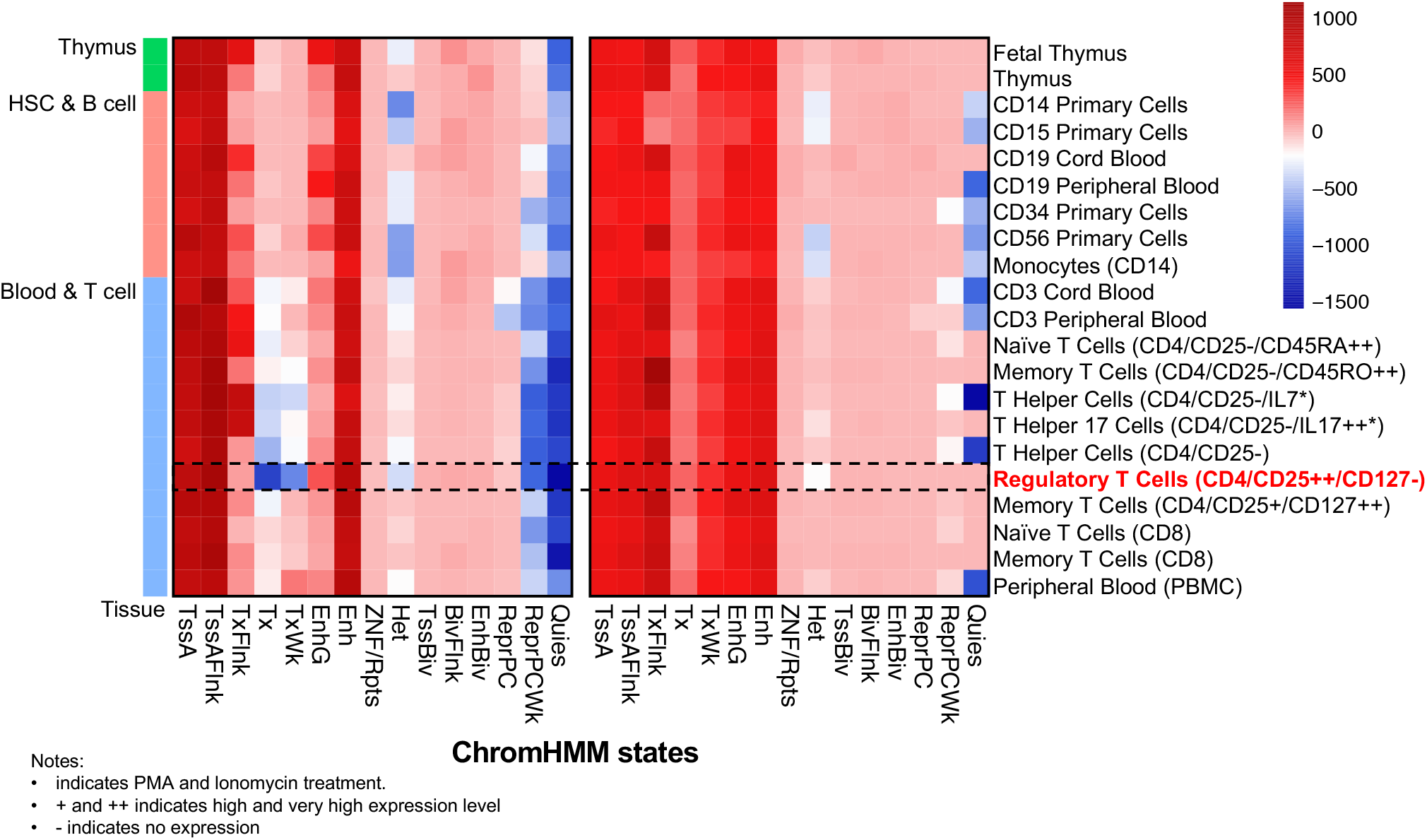
Enrichment of 3DFAACTS gnomAD variants (left panel) and their interacting regions (right panel) found within NIH Epigenomics Roadmap samples. Enrichment test of filtered gnomAD SNPs against chromHMM states from 129 tissues and cell types from Epigenomics Roadmap using GIGGLE (121). Red coloured regions indicate positive enrichment of variants within cell-types and chromHMM states, while blue coloured regions indicate negative enrichment. Here we subset to enrichment in three tissue groups, including thymus, HSC & B cell and Blood & T cell, enrichment result of all samples can be found in Additional file 1: Figure S18 & 19.

Treg Hi-C data was used to explore the FOXP3-associated regulatory networks that include these SNPs in a Treg. For the regions identified to interact with the 7,900 variants located in FOXP3 binding regions by Hi-C we observed a strong positive enrichment of regulatory states such as *TssA*, *TssAFlnk*, *Tx*, *Txwk*, *EnhG* and *Enh* in blood, HSC, B and T cells, supporting a regulatory role for these interacting regions (Figure 6 and Additional file 1: Figure S17). In total 3,245 Treg expressed genes (mean FPKM > 1) (65) were found to be associated by Hi-C with variants identified by 3DFAACTS-SNP analysis of the common SNP gnomAD dataset. GO and Hallmark genes sets from the Molecular Signatures Database (MSigDB) (123,124) analysis of these 3,245 interacting Treg expressed genes were significantly enriched (adjusted P-value < 0.05) in relevant GO terms such as T cell activation and regulation of hematopoiesis (Additional file 1: Figure S18) and autoimmune/Tregs-related gene sets, including TNFα via NF-κB, IL6/JAK/STAT3, and IL2/STAT5 signaling pathways (Additional file 1: Figure S19). Integration of the filtered gnomAD variants with *cis* Treg eQTLs from the DICE database (64), further identified 943 common variants previously demonstrated to impact gene expression in Tregs (Additional file 8: Table S9). These 943 variants are connected by Hi-C interactions to 1038 genes in our analysis of which 121 (11.6%) form a *cis* eQTL pair with the 3DFAACTS identified SNPs. Importantly, interacting genes were significantly enriched (Fisher exact test, P value = 9.06e-24) in genes that are associated with 49 autoimmune diseases from GAAD (54) supporting the idea that we have identified potential novel disease associated molecular mechanisms.

We then integrated SNPs identified by 3DFAACTS-SNPs with the active TFBS dataset identified from Tregs ATAC-seq data by *HINT-ATAC* (37) (Additional file 8: Table S9) to identify potential molecular mechanisms of action of these non-coding SNPs. We found 870 filtered SNPs are located within active binding sites of 521 TFs indicating that they may impact TF binding. Accounting for the requirement of Treg expression of the TF (65) or its differential expression in Tregs compared to effector T cells (125), the number of variants with the potential to alter TF binding in a Treg was reduced to 693 and 108 variants respectively (Additional file 8: Table S9). Of the variants that potentially impact the binding of a TF expressed in a Treg, 19 were found to be an eQTL with its interacting gene partner identified by Hi-C. Included in this list were genes previously associated with Treg stability and viability, specific Treg subsets and pathways known to influence Treg differentiation and function. This is consistent with 3DFAACTS-SNP identifying potential novel variants that contribute to a Treg defect in disease. For example, Treg IL23R and FAS expression is associated with Treg/Th17 imbalances in IBD and the chronic inflammatory disease, acute coronary syndrome (126,127) and here using 3DFAACTS-SNP we predict rs1324551 and rs72676067 may contribute to this altered expression by disrupting the binding of the transcription factors RBPJ and POU2F2 respectively. Other genes are up-regulated in specific Treg subsets, including TBCID4 (follicular regulatory T cells) (128), ACTA2 (Placental bed uterine Tregs and tumour-infiltrating Treg) (66,129), and POLR1A (cold-exposed Brown adipose tissue Treg) (130), suggesting the identified common variants could lead to functional defects in these specific Treg subsets. A third group of genes have been shown to regulate growth factor signaling pathways that are known to influence Treg differentiation and function (131–133). In particular, we have identified variants that alter expression of the genes involved in TGF-β signaling (SPTBN1, CDC7 and SLC35F2) and WNT signaling (SPTBN1 and MCC). For example, 19 SNPs are linked to the SPTBN1 gene by Hi-C in our analysis, eight of which are identified as eQTLs with SPTBN1 in Tregs, and of these three (rs10170646, rs4455200 and rs13386146) overlap and potentially disrupt the binding of the transcription factors BCL6, HES2 and BATF-Jun heterodimer respectively prioritising these potential causative variants linked to allele-specific expression (ASE) of SPTBN1 in Treg. However further investigation is required to establish if altered SPTBN1 caused by these variants may contribute to any disease in response to TGF-β and WNT signaling pathways. Together, these data indicate that the 3DFAACTS SNP pipeline in combination with the gnomAD database has the potential to annotate novel disease associated variants and their potential molecular mechanisms of action, many of which have not previously been investigated in GWAS studies.

## Discussion

GWAS and fine-mapping studies have identified over 50 candidate regions for T1D progression (11,13,134), however a broad understanding of the underlying disease mechanism has been difficult to elucidate without relevant functional information derived from cell-specific material. With the availability of whole genome annotation, we see that the majority of genetic risk lies in non-coding regions of the genome and is enriched in regulatory regions including promoters and enhancers. Traditionally, to understand how these variants may function they have been assigned to the nearest gene or genes within a defined linear distance. However this approach ignores the role of three-dimensional connectivity by which enhancers and repressors function to regulate transcription (135–137).

Recent approaches use statistical co-localization tests to link potential causal SNPs and quantitative trait loci (QTLs) to identify the genes regulated by GWAS loci (138). These methods require many samples in the correct cell type or physiological context and to date work best for local/*cis* QTLs, generally less than 1Mb in linear distance (135). An alternative approach used in this study and others (139,140) is to make use of chromosome conformation capture data to directly connect disease-associated regulatory regions to their target genes. As growing cellular and genomics evidence indicate that dysregulation of the Treg compartment contributes to autoimmune disease (46,141,142), we generated a cell type-specific 3D interaction profile in human regulatory T cells to establish an *in silico,* candidate loci reduction method to identify T1D-candidate regions that function in a Treg and the genes they affect. Open chromatin regions identified by ATAC-seq and regulatory regions identified by epigenetic marks such as histone H3K27ac can number in the tens of thousands in a specific cell type (65,143), we therefore initially focused on regulatory regions bound by the Treg-specific transcription factor FOXP3 given the essential role of FOXP3 in the Treg functional phenotype we hypothesized that candidate variants that are found within open, FOXP3-bound regions are likely to alter immunological tolerance. In addition as different autoimmune-diseases share genetic risk regions (59) we speculated that by identifying specific genetic variants that may contribute to T1D through the dysregulation of regulatory T cell functional fitness, this could be via mechanisms consistent across many autoimmune diseases (1,144,145).

The design and implementation of the 3DFAACTS-SNPs workflow champions a new data-centric view of functional genomics analysis, with the development of cell type-specific epigenomic and 3D datasets enabling researchers to narrow down on molecular changes at a fine-scale resolution. However, results shown in this study suggests that cell type-specific viewpoints can be broadened to a much more lineage (T cell) or immune (e.g. innate or adaptive) system-specific level. While we focused on Treg cells and expected to identify Treg-specific enhancer-controlled targets, based on the criteria of inclusion of FOXP3 binding data, no functional variant was uniquely accessible in only Tregs, nor were they specifically enriched with Treg-exclusive H3K27ac modified regions (Figure 4B). This likely reflects the propensity of FOXP3 to bind to enhancers active in multiple CD4+ T cell lineages (78) (Figure 4) to modify their output in a Treg-specific manner and therefore we cannot currently discern whether these filtered variants act predominantly in Tregs or on other CD4+ T cell subsets. The incorporation of context- and CD4+ T cell subset-specific gene expression (146) and epigenomic (140,147) data into the 3DFAACTS-SNPs workflow may help resolve this. Although we have focused here on using FOXP3-binding as a filtering criteria, it is known that other FOXP3-independent pathways are important for Treg function and the 3DFAACTS-SNPs workflow could be modified to incorporate other TFs or other epigenetic profiles such as CpG-demethylated regions (148) to further explore the relationship between disease-associated variants and these pathways.

In total using the 3DFAACTS-SNPs workflow we identified 36 novel candidate genes connected to variants in 12 T1D risk loci that could plausibly function in a Treg whereas we could not define plausible candidate Treg-specific activity at the other T1D risk regions that met all our filtering criteria. This may indicate that these other risk-regions are active in immune cell types other than a Treg or they impact genes and regulatory elements within a Treg that are not dependent upon FOXP3. As an example of how the 3DFAACTS-SNPs workflow can lead to testable insights into the molecular mechanisms of non-coding variants, the SNP rs614120 was found to be located in a FANTOM5 annotated T cell-specific enhancer region in the first intron of the BACH2 gene, and is predicted to disrupt the binding of Forkhead Transcription factor family member FOXA2 (Figure 5 and Additional file 6: Table S7). However, FOXA2 is not expressed in T cells, indicating that rs614120 might disrupt the binding of other Forkhead family members which bind to very similar DNA sequences, such as FOXP3, which is known to bind in this region (Figure 5). The 3DFAACTS-SNPs workflow further indicates that this enhancer region containing rs614120 interacts with the promoter of BACH2, forming a distal promoter-enhancer interactions, suggesting that rs614120 may disrupt FOXP3 binding to the enhancer leading to the dysregulation of BACH2 expression. It has been recently shown that Bach2 plays roles in the regulation of T cell receptor signalling in Tregs, including averting premature differentiation and assisting peripherally induced Treg development (149). Therefore, we suggested that this single variant may regulate BACH2 expression and ultimately may affect the progression of T1D, and this requires further experiments to verify. This can further aid the development of novel therapeutic approaches to restore function in Treg of patients with this genotype. This finding also suggests that variants can contribute to the causal mechanisms of disease by altering the efficacy/ stability of TF binding in important regions such as enhancers or SEs.

The power of 3DFAACTS-SNPs is its ability to incorporate chromosome organisation in 3D and identify long-range interactions involving variant-containing regulatory regions leading to the identification of target genes that have not previously been associated with these disease associated risk regions. This is illustrated by the finding that the majority (24/31) of Treg-expressed genes that interact with the T1D variants are not the closest gene in linear proximity and of these interacting genes 20 have not been previously associated with any autoimmune disease. For example, T1D 3DFAACTS SNP rs1029991 although located in linear proximity to the CD69 gene was found to contact the CHD4 gene (∼3.2 mb away) (Additional file 3: Table S4) suggesting this variant is more likely to influence CHD4 expression than CD69. Interestingly, rs1029991 was not identified as a *cis*-eQTL for CHD4 in Tregs as it >3Mb away on the genome, with eQTLs being classified as cis when found <1Mb from their target gene.

The idea that high-order nuclear organisation coordinates transcription in times of immune challenge or tolerance was recently shown in a study demonstrating that 3D chromatin looping topology is important for a subset of long non-coding RNAs (lncRNAs), termed immune gene– priming lncRNAs (IPLs), to be correctly positioned at the promoters of innate genes (70). This positioning of the IPLs then allows for the recruitment of the WDR5–mixed lineage leukaemia protein 1 (MLL1) complex to these promoters to facilitate their H3K4me3 epigenetic priming (70). An example of long-range enhancer gene interactions in conveying autoimmune-disease risk in Treg cells has also recently been published (150). In this work a distal enhancer at the 11q13.5 locus associated with multiple autoimmune-disease risk, including T1D was found to participate in long-range interactions with the LRRC32 gene exclusively in Treg. Deletion of this enhancer in mice resulted in the specific loss of Lrcc32 expression in Treg cells and the inability of Treg to control gut-inflammation in an adoptive transfer colitis model. Furthermore CRISPR-activation experiments in human Tregs identified a regulatory element located in proximity to a risk variant rs11236797 that is capable of influencing LRRC32 expression. This data together highlights the mechanistic basis of how non-coding variants may function to interfere with Treg activity in disease. Although we did not identify this interaction in our final SNP-interaction list upon re-examination of our workflow this interaction was present in our Hi-C dataset, but it was filtered out as the enhancer is not bound by FOXP3. Coordinated genome topology has also been shown in immune cell lineage commitment, both at a loci (151,152) and compartment level (153), consistent with the concept of immune transcriptional “factories” where genes congregate in regions of the nucleus to undergo coordinated transcriptional activation (154).

Although a shared genetic aetiology between T1D and other immune-mediated diseases has been proposed we did not find a large overlap between the variants or interacting genes identified by 3DFAACTS SNP in T1D and other autoimmune disease datasets. The reason for this is not clear but may be a result of the relatively low number of input SNPs for the other autoimmune diseases. Irrespective of this, several candidate causal SNPs and genes including SFMBT2 (rs61839660), ABI2 and IQCA1 (rs3087243) and PFKFB3 (rs12722496 and rs12722508 - T1D and rs947474 - 4AI) were found to be common between T1D and other autoimmune diseases. Several of these genes such as SFMBT2, ABI2 and PFKFB3 have previously been implicated in the development of autoimmune diseases or play a role in critical T cell pathways suggesting these genes are likely targets that explain the molecular function of the risk variants. SFMBT2 is a methylated histone binding transcriptional repressor which has been associated with childhood onset asthma (155). ABI2 is required for actin polymerization at the T cell:APC contact site with loss of Abi1 in mice resulting in decreased TCR-mediated IL-2 production and proliferation (156). PFKFB3 is involved in both the synthesis and degradation of fructose-2,6-bisphosphate, a regulatory molecule that controls glycolysis in eukaryotes. Regulation of glycolysis has increasingly been implicated in shaping immune responses (157) and PFKFB3 has been associated with multiple autoimmune diseases (158). Importantly, reduced PFKFB3 enzyme activity leading to redox imbalance and apoptosis has been reported in CD4+ T from RA patients (159) directly linking the PFKFB3 gene to the disease.

A highly polygenic architecture with small effect sizes of many causal variants (112,113) has been proposed to account for missing heritability associated with phenotypic traits. Most of these small effect size variants have yet to be identified. Here we have begun to investigate whether common genetic variation found within populations could contribute to autoimmune diseases by altering gene-expression by altering enhancer and promoter output. In this study we illustrate this potential by accessing large population-scale variant resources in the gnomAD database, identifying 7,900 filtered common variants that have the potential to impact Treg function. Based on the search of discovered associations of autoimmune diseases (EFO_0005140) from the GWAS Catalog (160), over half of the variants surveyed here have not been used in large-scale autoimmune disease GWAS (11,93,161–167), precluding their assessment for potential disease risk in sampled disease/control populations. While filtered variants identified here are biased towards the inclusion of FOXP3-binding within the workflow, their potential immune response impact is highlighted by the finding that their interacting regions are positively enriched for transcription and enhancer-associated chromatin states (Figure 6, Additional file 1: Figure S16 & 17), eQTLs and potentially impacted TFBS (Additional file 8: Table S9). This potential accessibility of regulatory variants among a population could potentially explain additional variation in effector responses in T cell activation (168), relevant not only to autoimmune disease, but also to broader immune responses for example to SARS-CoV-2.

In conclusion, while we initially restricted the application of 3DFAACTS-SNP to Treg centric genome-wide interaction frequency profiles to give functional annotation in T1D data, we have demonstrated that valid interacting pairs from Hi-C dataset can be functionally mapped with high confidence from multiple disease datasets as well as whole genome variant datasets, which presents a valuable resource in establishing cell-type specific interactomes. Coupled with cell-type specific genomic data available from public repositories, such as the NIH Roadmap (52), Blueprint (169) and ENCODE (170) projects, this workflow provides a useful mechanism to identify potential mechanisms by which non-coding variants regulate disease causing genes, and identifies new targets for therapeutic modulation to treat or prevent disease.

## Conclusion

Based on Treg ATAC-seq, Hi-C data, promoters and enhancers annotation and FOXP3 binding site, we developed a variant filtering workflow named 3DFAACTS-SNP to identify potential causative SNPs and their 3D interacting genes for T1D from GWAS fine-mapped variants. Our workflow can easily be used with variants associated with other autoimmune diseases or even large population-scale variants.

## List of abbreviations

GWAS: Genome-wide association study
SNP: Single nucleotide polymorphism
T1D: Type 1 diabetes
Treg: Regulatory T cells
Tconv: Conventional T cells
FOXP3: Foxhead box protein 3
TF: Transcription factor
eQTL: Expression quantitative trait loci
ChIP-seq: Chromatin immunoprecipitation sequencing
ATAC-seq: Assay for transposase-accessible chromatin sequencing
Hi-C: High resolution chromosome conformation capture sequencing
TAD: Topologically-associated domain
SE: Super-enhancer

## Declarations

### Ethics approval and consent to participate

Cord blood used in this study was obtained with approval from the donor and the Children’s, Youth and Women’s Health Service Research Ethics Committee (HREC1596 and HREC 19/wchn/65 from the Women’s and Children’s Health Network Human Ethics committee). Buffy Coats were obtained from the Australian Red Cross (Material Supply Deed 19-03SA-02).

### Consent for publication

Not applicable

### Availability of data and materials

The Treg ATAC-seq and Hi-C datasets analysed during the current study are available in the European Nucleotide Archive (ENA) repository (PRJEB39882).

#### Published data

FOXP3 ChIP-chip data used during the current study is available on Gene Expression Omnibus (accession no. GSE20995) (47).

#### Database used in the current study

NIH Roadmap Epigenomics Project: http://www.roadmapepigenomics.org/ (52)

SEdb: http://www.licpathway.net/sedb/ (76)

GAAD: http://gaad.medgenius.info/intro/ (54)

DICE: https://dice-database.org/landing (64)

gnomAD: https://gnomad.broadinstitute.org/ (120)

FANTOM5: https://fantom.gsc.riken.jp/5/ (56)

Source code for 3DFAACTS-SNP workflow and related in-house scripts are available in GitHub (https://github.com/ningbioinfostruggling/3DFAACTS-SNP).

### Authors information

Ning Liu and Timothy Sadlon are joint first authors. Simon Barry and James Breen are joint corresponding authors

### Contributions

All authors designed the study design. NL, TS, SB and JB wrote the manuscript, with additional help from YYW and SMP. NL developed and implemented all computational workflows and HiC-seq analyses. YYW analysed the ATAC-seq datasets and JB analysed public ChIP-seq and ATAC-seq data. TS and YYW constructed the ATAC-seq and Hi-C libraries.

## Acknowledgements

We are thankful for the access to publicly available resources, and grateful for discussions and comments from members of the Barry lab. We thank Dr R Gross for cell sorting expertise.

## Supplementary information

### Additional file 1

Figures. S1–19, and Tables S1-2.

### Additional file 2: Table S3

Promoters and enhancers used in the 3DFAACTS-SNP workflow in this study.

### Additional file 3: Table S4

T1D 3DFAACTS SNPs identified using 3DFAACTS-SNP workflow from T1D fine-mapped SNPs and their 3D interacting genes.

### Additional file 4: Table S5

Topologically-associated domains (TADs) identified using TopDom from Treg Hi-C data and the relationship between Hi-C interactions of T1D 3DFAACTS SNPs and identified TADs.

### Additional file 5: Table S6

Transcription factor footprint identified from rest and stimulated Tregs ATAC-seq data using HINT-ATAC.

### Additional file 6: Table S7

T1D 3DFAACTS SNPs that are located in active transcription factor footprint in Tregs. And the binding affinity effect of these SNPs with their overlapped transcription factor calculated using GWAS4D.

### Additional file 7: Table S8

3DFAACTS SNPs identified from fine-mapped and meta-analysis SNP datasets from 3 published studies.

### Additional file 8: Table S9

3DFAACTS SNPs identified from gnomAD common (MAF >= 0.1) SNPs.

