## Additional File 1 for "3DFAACTS-SNP: Using regulatory T cell-specific epigenomics data to uncover candidate mechanisms of Type-1 Diabetes (T1D) risk"

**Supplementary Tables**

| Metrics | Rest Treg D1 | Stim Treg D1 | Rest Treg D2 | Stim Treg D2 | Rest Treg D3 | Stim Treg D3 |
| --- | --- | --- | --- | --- | --- | --- |
| Raw reads | 41,157,156 | 42,588,662 | 32,775,125 | 38,930,881 | 34,870,964 | 32,230,708 |
| Mapped reads | 36,592,827 (88.91%) | 39,454,136 (92.64%) | 29,327,182 (89.48%) | 35,092,296 (90.14%) | 31,307,151 (89.78%) | 29,726,382 (92.23%) |
| Uniquely mapped | 27,970,773 (67.96%) | 32,704,650 (76.79%) | 22,653,938 (69.12%) | 28,629,417 (73.54%) | 24,474,978 (70.19%) | 24,870,158 (77.16%) |
| Duplication | 3.90% | 6.60% | 4.20% | 6.70% | 4% | 7.20% |

**Table S1.** Statistics of Tregs ATAC-seq data.

| **Metrics** | **Tregs Hi-C rep1** | **Tregs Hi-C rep2** |
| --- | --- | --- |
| Sequenced Read Pairs | 3,020,186 | 2,678,685 |
| Unmapped Reads | 196,702 (6.513%) | 139,223 (5.197%) |
| Unique Aligned Pairs | 1,973,216 (65.334% / 100%) | 1,715,393 (64.039% / 100%) |
| Valid Contacts | 459,244 (15.21% / 23.27%) | 1,441,362 (53.81% / 84.03%) |
| Duplicate Contacts | 5,525 (0.18% / 0.28%) | 16,872 (0.63% / 0.98%) |
| Inter Chromosomal | 128,073 (4.24% / 6.49%) | 345,409 (12.89% / 20.14%) |
| Intra Chromosomal | 325,646 (10.78% / 16.5%) | 1,079,081 (40.28% / 62.91%) |
| Intra Short Range (< 20kb) | 88,121 (2.92% / 4.47%) | 243,166 (9.08% / 14.18%) |
| Intra Long Range (> 20kb) | 237,525 (7.86% / 12.04%) | 835,915 (31.21% / 48.73%) |
| Read Pair Type (L-I-O-R) | 24.71%-26.32%-24.44%-24.53% | 25.05%-25.23%-24.72%-25.0% |

**Table S2.** Statistics of Tregs Hi-C data.

**Table S3**: Promoters and enhancers used in the 3DFAACTS-SNP workflow in this study.

Table S3 is shown in a separate file (Additional File 2).

**Table S4**: T1D 3DFAACTS SNPs identified using 3DFAACTS-SNP workflow from T1D fine-mapped SNPs and their 3D interacting genes.

Table S4 is shown in a separate file (Additional File 3).

**Table S5**: Topologically-associated domains (TADs) identified using TopDom from Treg Hi-C data and the relationship between Hi-C interactions of T1D 3DFAACTS SNPs and identified TADs.

Table S5 is shown in a separate file (Additional File 4).

**Table S6**: Transcription factor footprint identified from rest and stimulated Tregs ATAC-seq data using HINT-ATAC.

Table S6 is shown in a separate file (Additional File 5).

**Table S7**: T1D 3DFAACTS SNPs that are located in active transcription factor footprint in Tregs. And the binding affinity effect of these SNPs with their overlapped transcription factor calculated using GWAS4D.

Table S7 is shown in a separate file (Additional File 6).

**Table S8**: 3DFAACTS SNPs identified from fine-mapped and meta-analysis SNP datasets from 3 published studies.

Table S8 is shown in a separate file (Additional File 7).

**Table S9**: 3DFAACTS SNPs identified from gnomAD common (MAF >= 0.1) SNPs.

Table S9 is shown in a separate file (Additional File 8).

**Supplementary Figures**

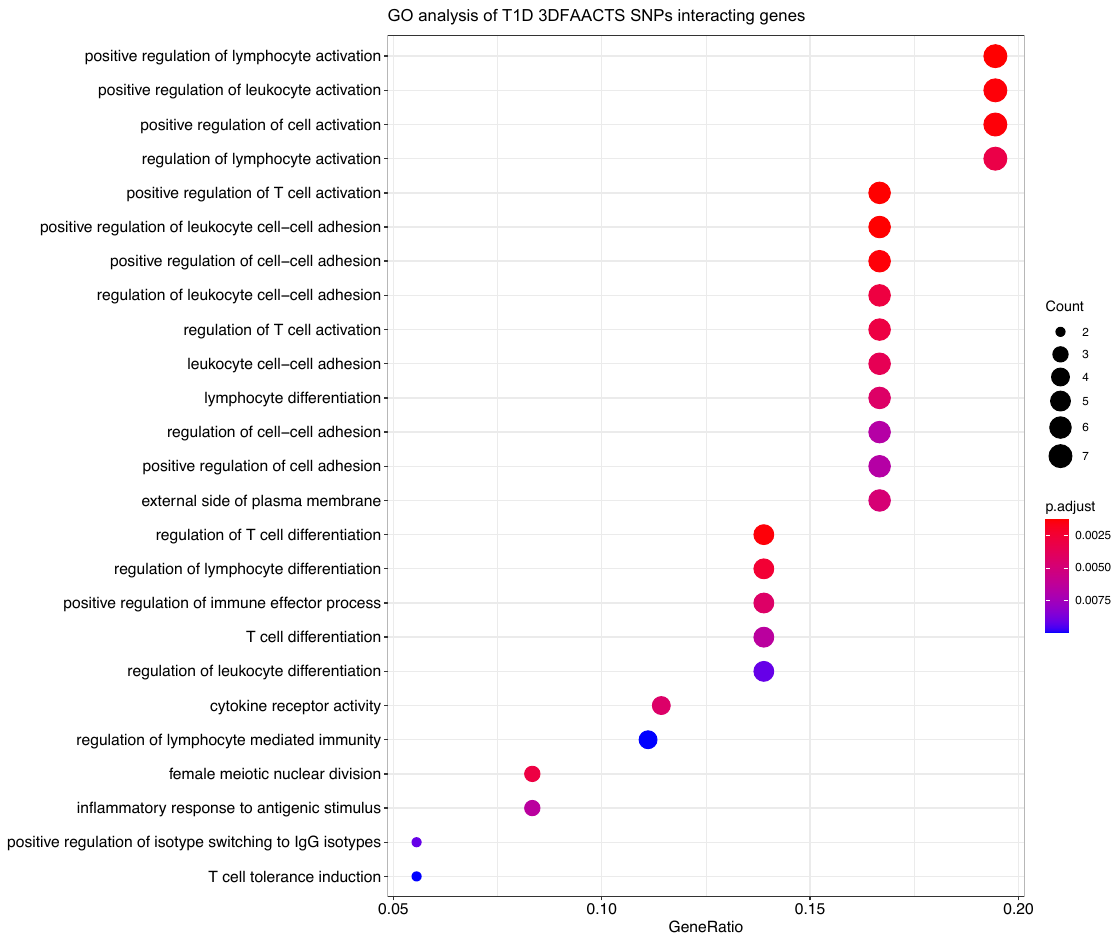

**Figure S1: Top20 significantly enriched gene ontology (GO) terms of T1D 3DFAACTS SNPs interacting genes.**

Color indicates adjusted P-value of the enrichment test, and the sizes of the dots indicate the number of genes are included in the GO term.

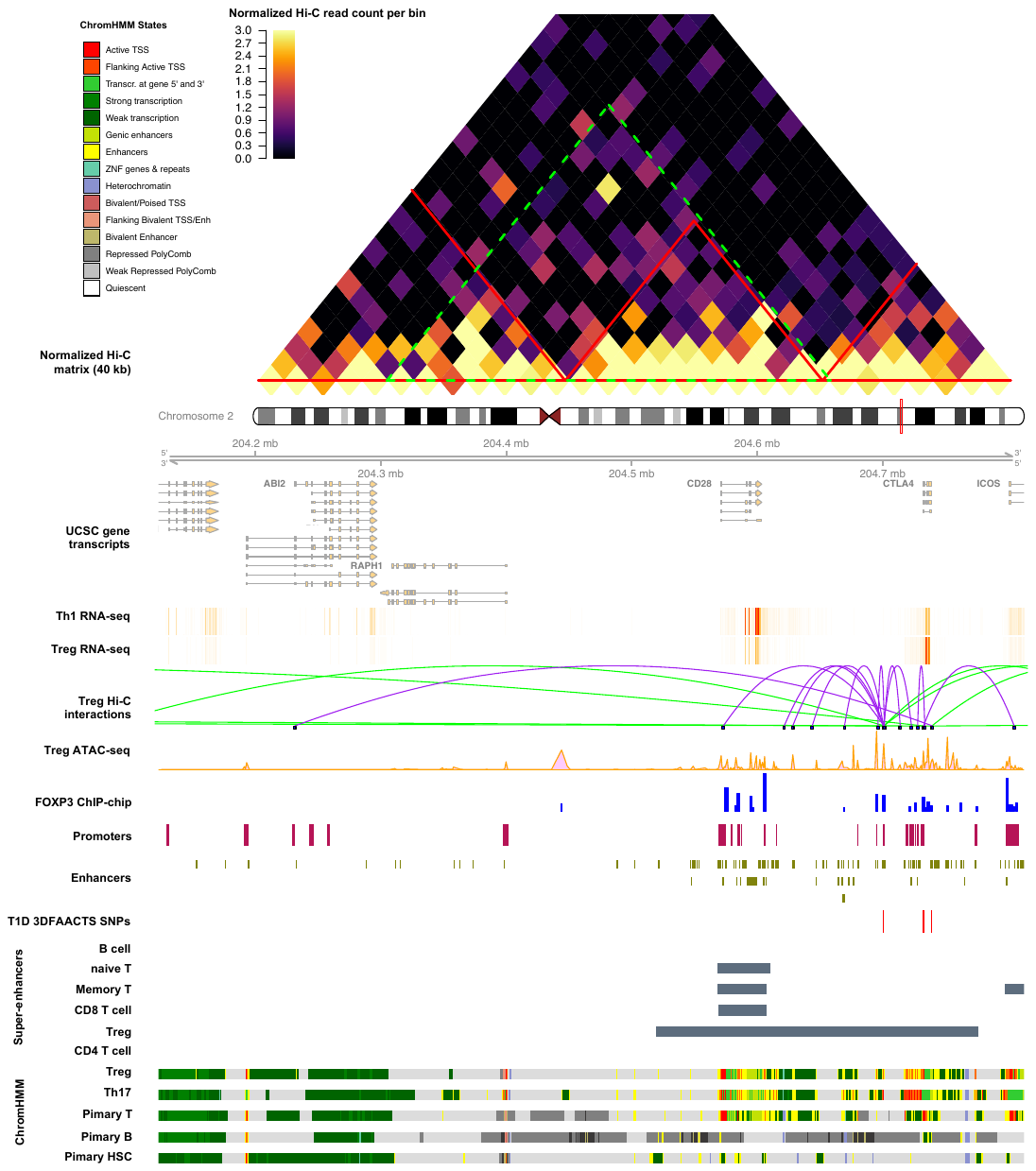

**Figure S2:** **Visualisation of the region of filtered T1D SNPs on chromosome 2.** Heatmap shows the Tregs Hi-C normalised interaction matrix (resolution of 40kb) on chr2: 203922714-205092714. The red triangles indicate Topologically Associated Domains (TADs) and the large green-dotted triangle indicates the boundary of the current plot. Tracks displayed below the chromosome 2 ideogram display workflow datasets (filtered SNPs, FOXP3-binding sites and Treg ATAC-seq, Hi-C interactions, promoters and enhancers) along with various types of cell type-specific data including UCSC Gene Transcript information, T cell subsets (Thelper1 and Treg) expression data, super-enhancer data and 15-state ChromHMM track of T cell lineages.

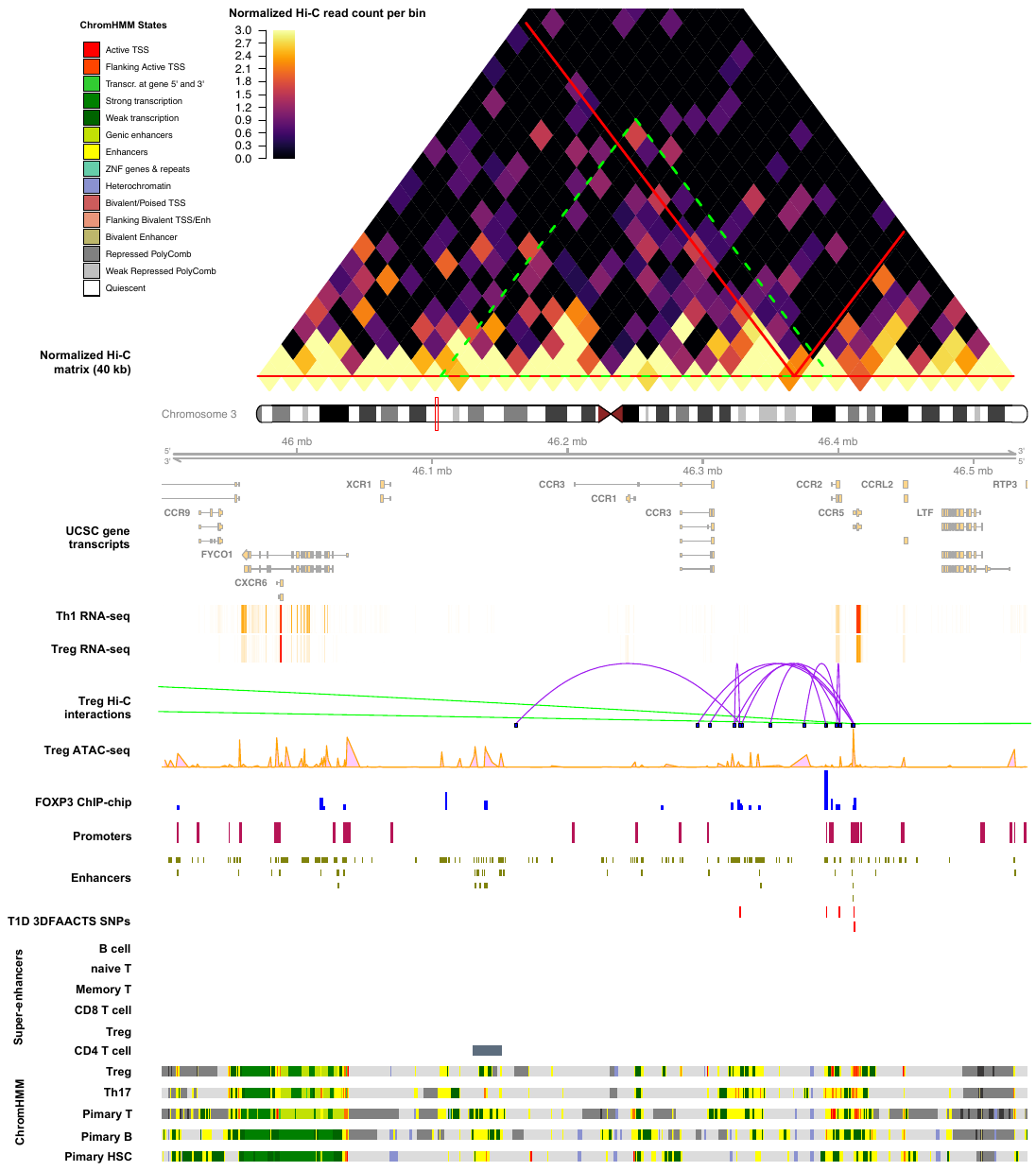

**Figure S3:** **Visualisation of the region of filtered T1D SNPs on chromosome 3.** Heatmap shows the Tregs Hi-C normalised interaction matrix (resolution of 40kb) on chr3: 45600000-46840000. The red triangles indicate Topologically Associated Domains (TADs) and the large green-dotted triangle indicates the boundary of the current plot. Tracks displayed below the chromosome 3 ideogram display workflow datasets (filtered SNPs, FOXP3-binding sites and Treg ATAC-seq, Hi-C interactions, promoters and enhancers) along with various types of cell type-specific data including UCSC Gene Transcript information, T cell subsets (Thelper1 and Treg) expression data, super-enhancer data and 15-state ChromHMM track of T cell lineages.

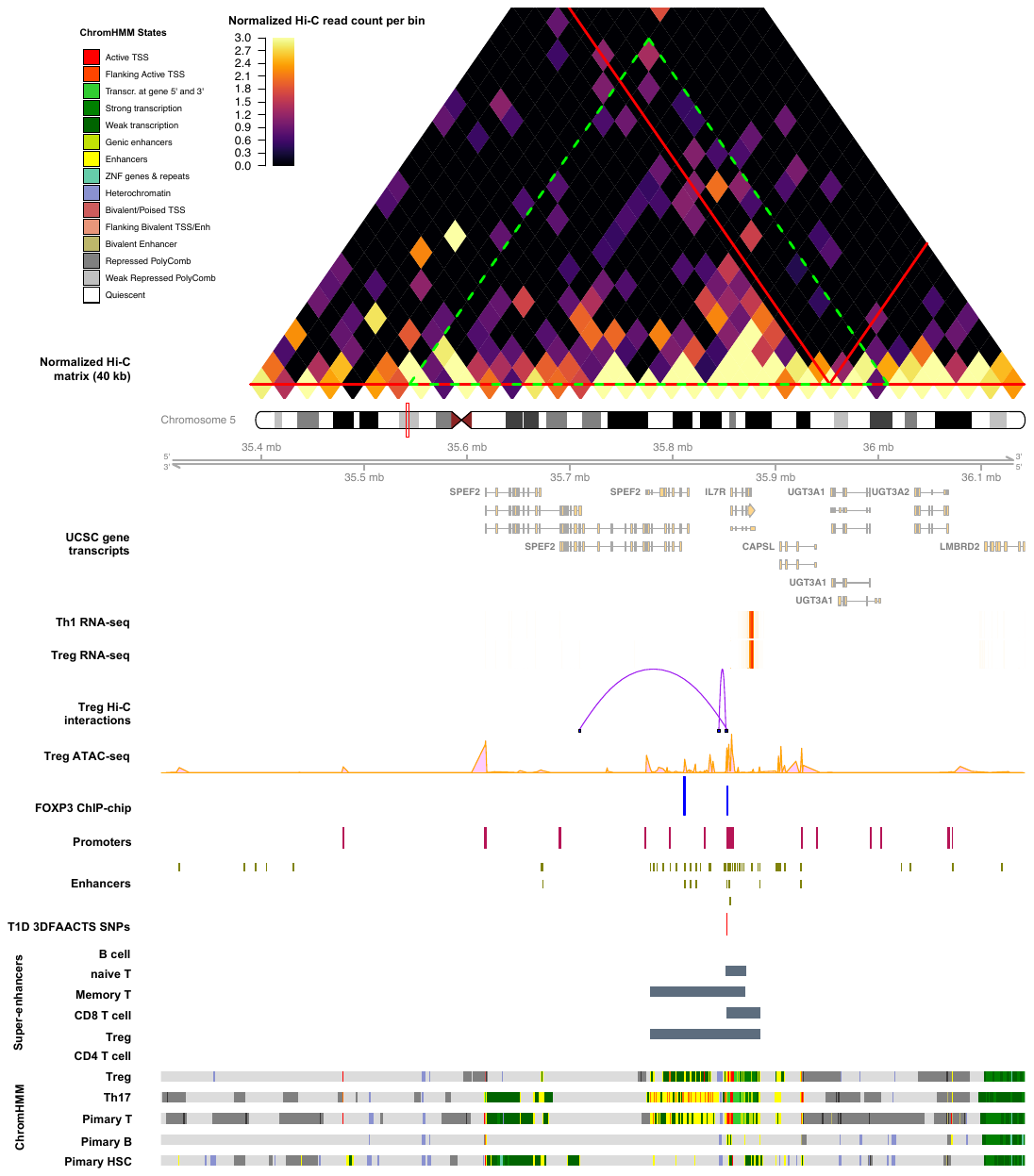

**Figure S4:** **Visualisation of the region of filtered T1D SNPs on chromosome 5.** Heatmap shows the Tregs Hi-C normalised interaction matrix (resolution of 40kb) on chr5: 35022311-36382311. The red triangles indicate Topologically Associated Domains (TADs) and the large green-dotted triangle indicates the boundary of the current plot. Tracks displayed below the chromosome 5 ideogram display workflow datasets (filtered SNPs, FOXP3-binding sites and Treg ATAC-seq, Hi-C interactions, promoters and enhancers) along with various types of cell type-specific data including UCSC Gene Transcript information, T cell subsets (Thelper1 and Treg) expression data, super-enhancer data and 15-state ChromHMM track of T cell lineages.

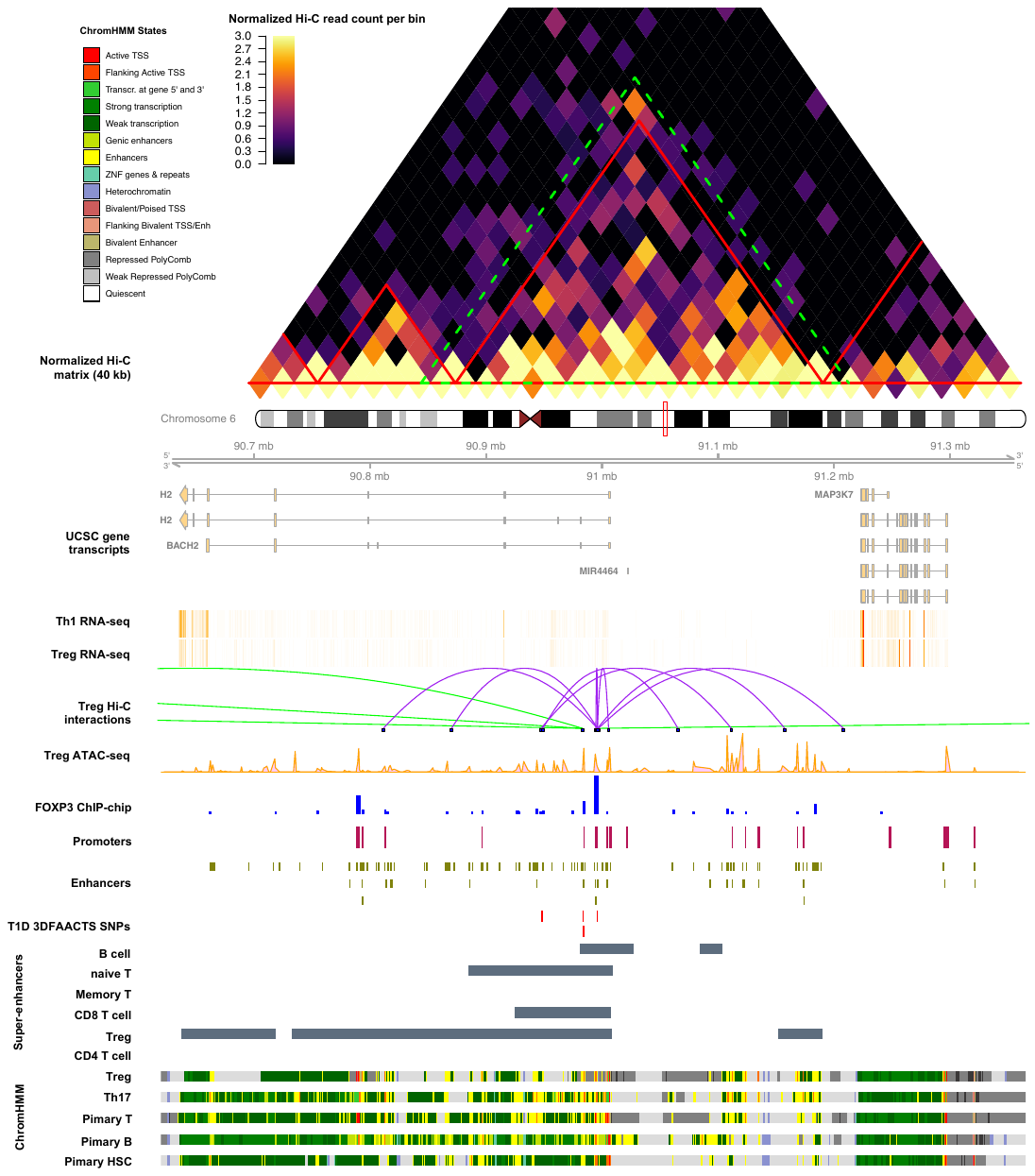

**Figure S5:** **Visualisation of the region of filtered T1D SNPs on chromosome 6.** Heatmap shows the Tregs Hi-C normalised interaction matrix (resolution of 40kb) on chr6: 90320000-91665000. The red triangles indicate Topologically Associated Domains (TADs) and the large green-dotted triangle indicates the boundary of the current plot. Tracks displayed below the chromosome 6 ideogram display workflow datasets (filtered SNPs, FOXP3-binding sites and Treg ATAC-seq, Hi-C interactions, promoters and enhancers) along with various types of cell type-specific data including UCSC Gene Transcript information, T cell subsets (Thelper1 and Treg) expression data, super-enhancer data and 15-state ChromHMM track of T cell lineages.

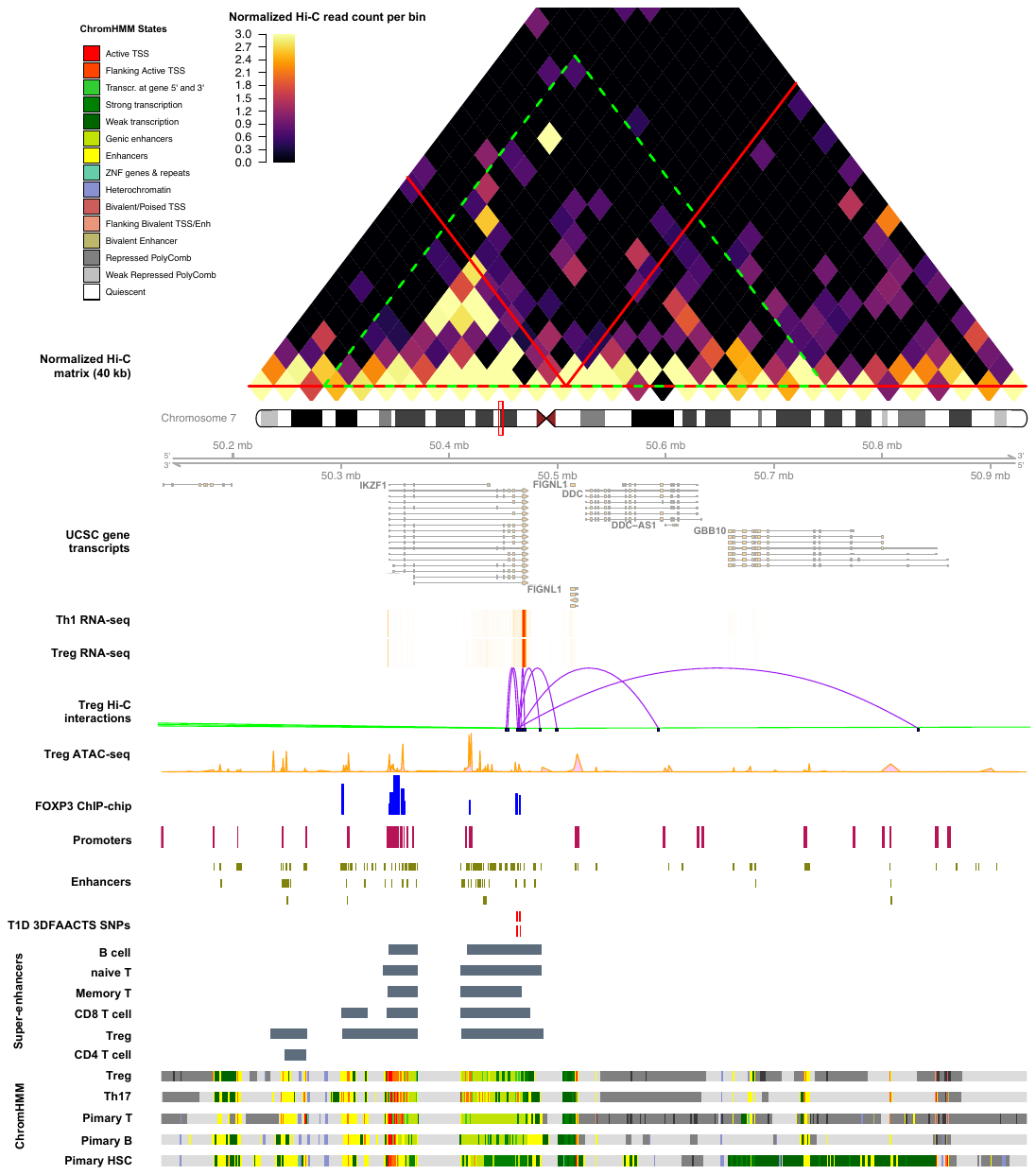

**Figure S6:** **Visualisation of the region of filtered T1D SNPs on chromosome 7.** Heatmap shows the Tregs Hi-C normalised interaction matrix (resolution of 40kb) on chr7: 50013852-51253852. The red triangles indicate Topologically Associated Domains (TADs) and the large green-dotted triangle indicates the boundary of the current plot. Tracks displayed below the chromosome 7 ideogram display workflow datasets (filtered SNPs, FOXP3-binding sites and Treg ATAC-seq, Hi-C interactions, promoters and enhancers) along with various types of cell type-specific data including UCSC Gene Transcript information, T cell subsets (Thelper1 and Treg) expression data, super-enhancer data and 15-state ChromHMM track of T cell lineages.

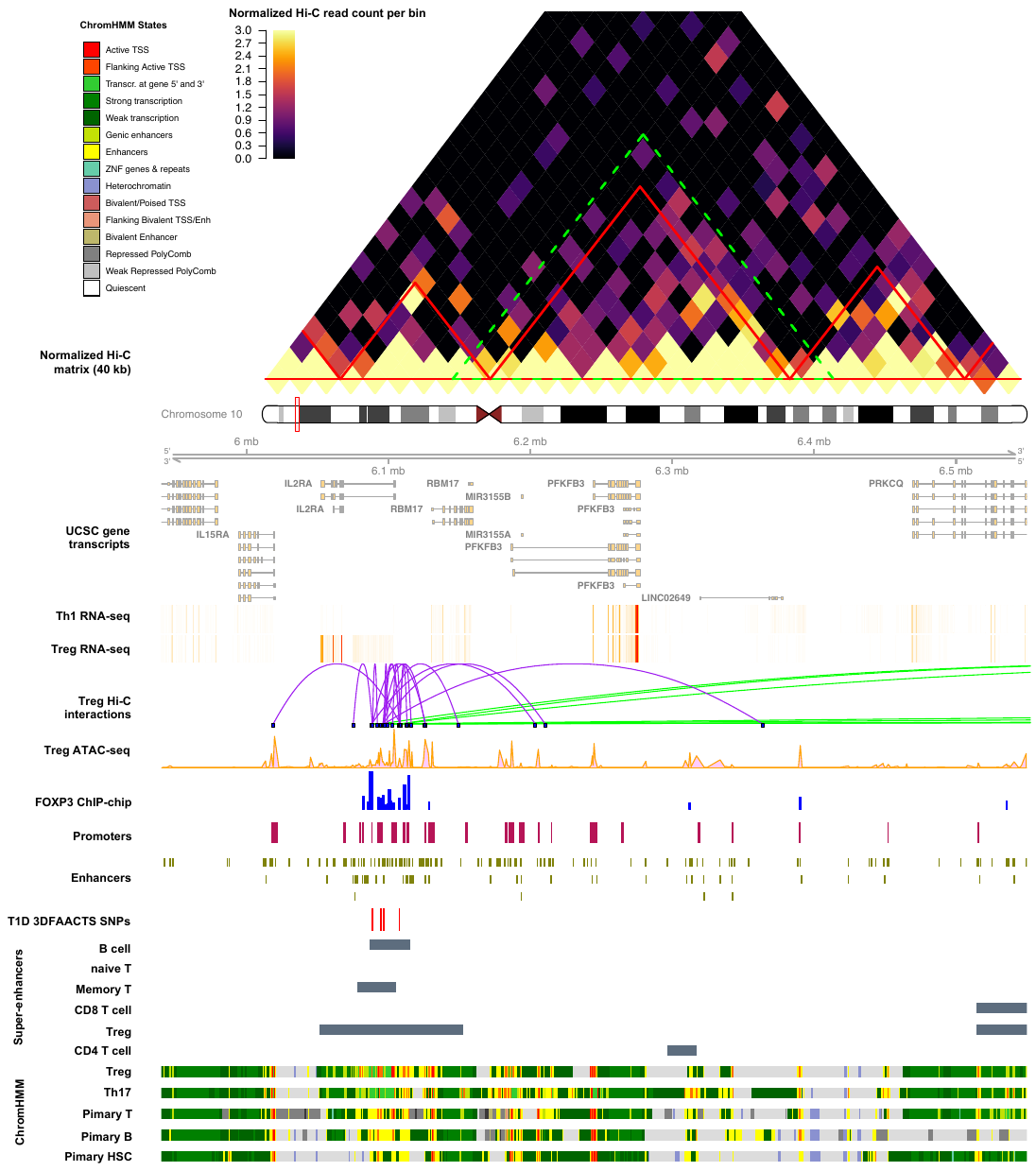

**Figure S7:** **Visualisation of the region of filtered T1D SNPs on chromosome 10.** Heatmap shows the Tregs Hi-C normalised interaction matrix (resolution of 40kb) on chr10: 5640000-6850000. The red triangles indicate Topologically Associated Domains (TADs) and the large green-dotted triangle indicates the boundary of the current plot. Tracks displayed below the chromosome 10 ideogram display workflow datasets (filtered SNPs, FOXP3-binding sites and Treg ATAC-seq, Hi-C interactions, promoters and enhancers) along with various types of cell type-specific data including UCSC Gene Transcript information, T cell subsets (Thelper1 and Treg) expression data, super-enhancer data and 15-state ChromHMM track of T cell lineages.

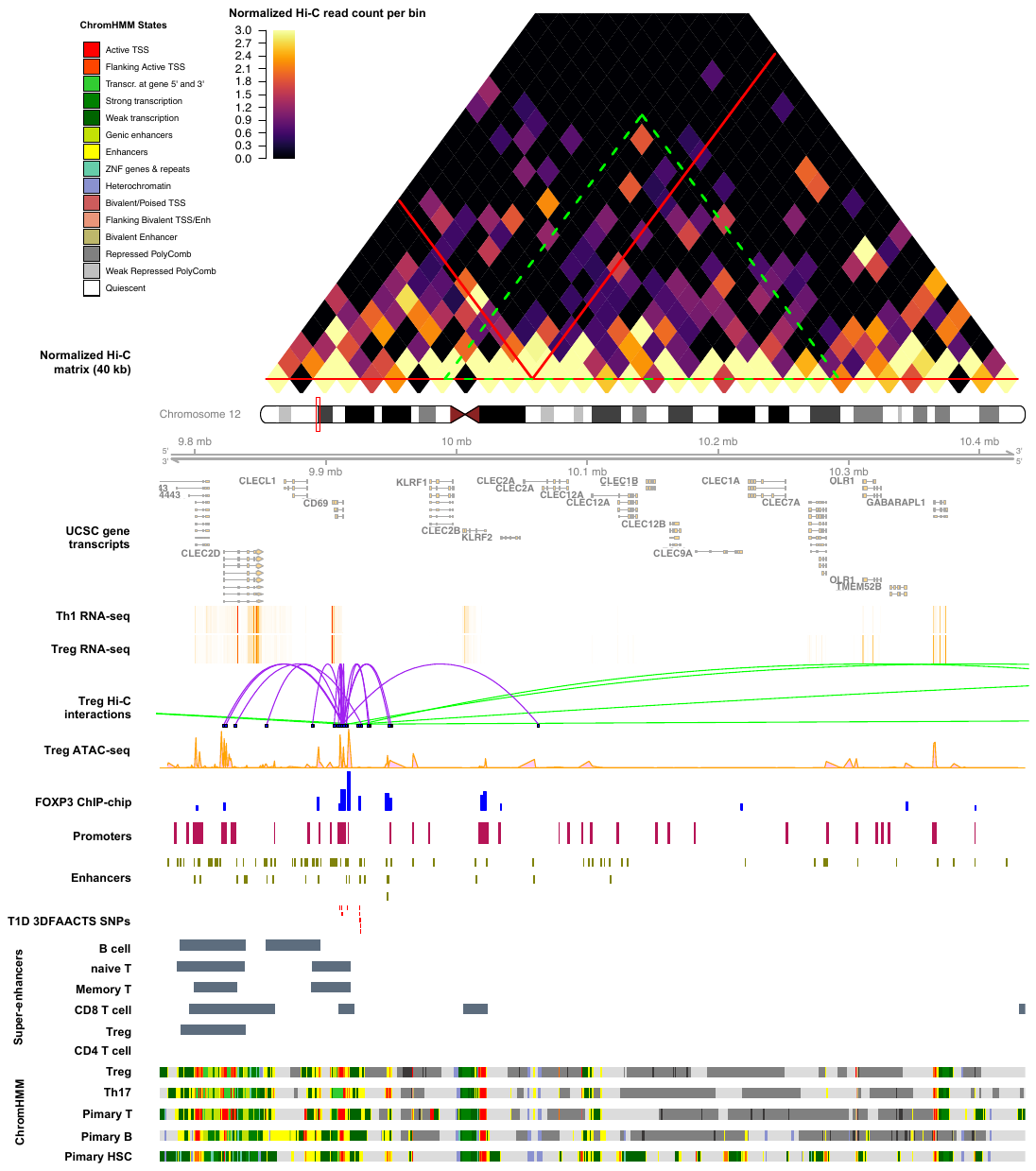

**Figure S8:** **Visualisation of the region of filtered T1D SNPs on chromosome 12.** Heatmap shows the Tregs Hi-C normalised interaction matrix (resolution of 40kb) on chr12: 9473000-10735000. The red triangles indicate Topologically Associated Domains (TADs) and the large green-dotted triangle indicates the boundary of the current plot. Tracks displayed below the chromosome 12 ideogram display workflow datasets (filtered SNPs, FOXP3-binding sites and Treg ATAC-seq, Hi-C interactions, promoters and enhancers) along with various types of cell type-specific data including UCSC Gene Transcript information, T cell subsets (Thelper1 and Treg) expression data, super-enhancer data and 15-state ChromHMM track of T cell lineages.

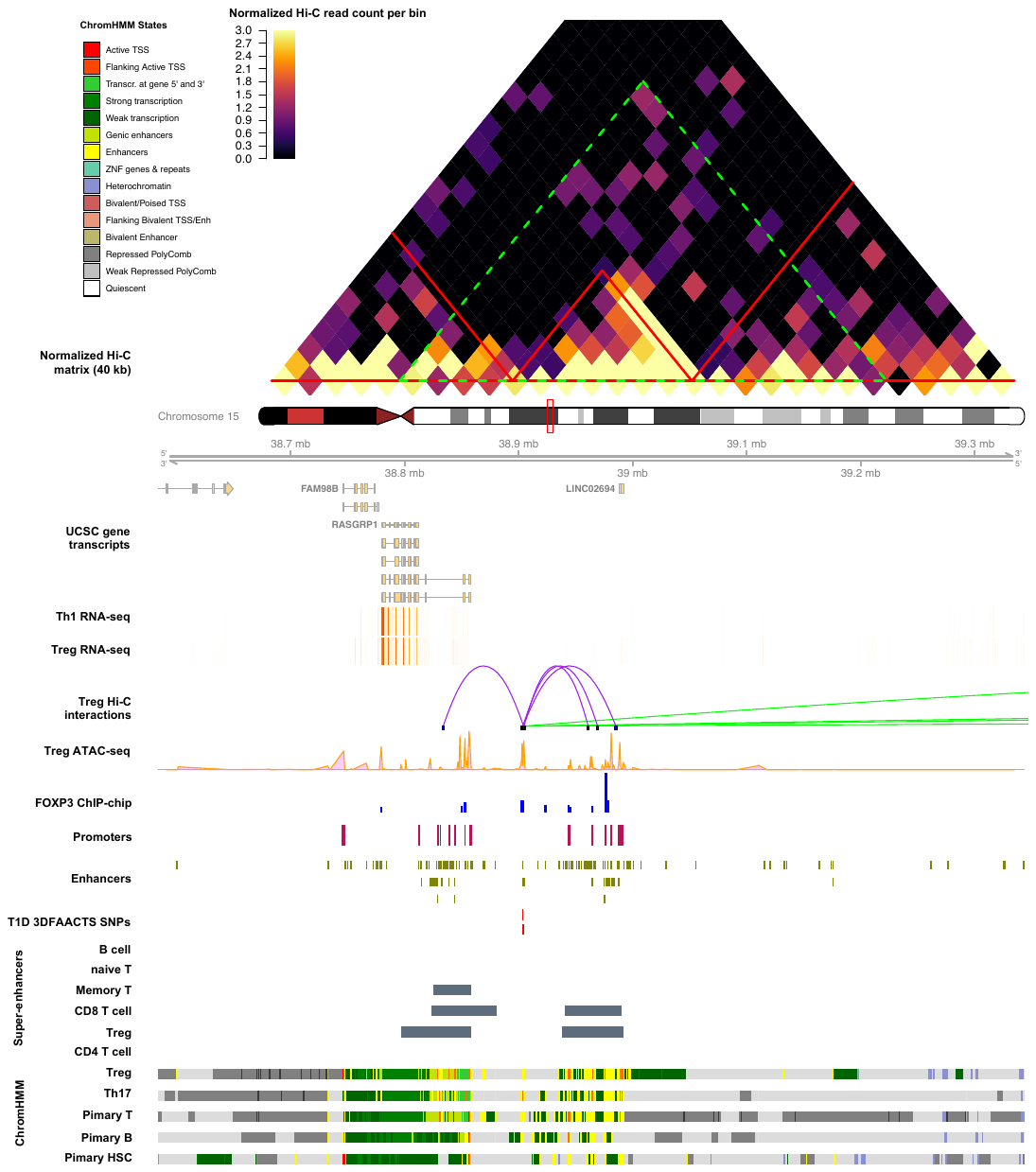

**Figure S9:** **Visualisation of the region of filtered T1D SNPs on chromosome 15.** Heatmap shows the Tregs Hi-C normalised interaction matrix (resolution of 40kb) on chr15: 38383672-39543672. The red triangles indicate Topologically Associated Domains (TADs) and the large green-dotted triangle indicates the boundary of the current plot. Tracks displayed below the chromosome 15 ideogram display workflow datasets (filtered SNPs, FOXP3-binding sites and Treg ATAC-seq, Hi-C interactions, promoters and enhancers) along with various types of cell type-specific data including UCSC Gene Transcript information, T cell subsets (Thelper1 and Treg) expression data, super-enhancer data and 15-state ChromHMM track of T cell lineages.

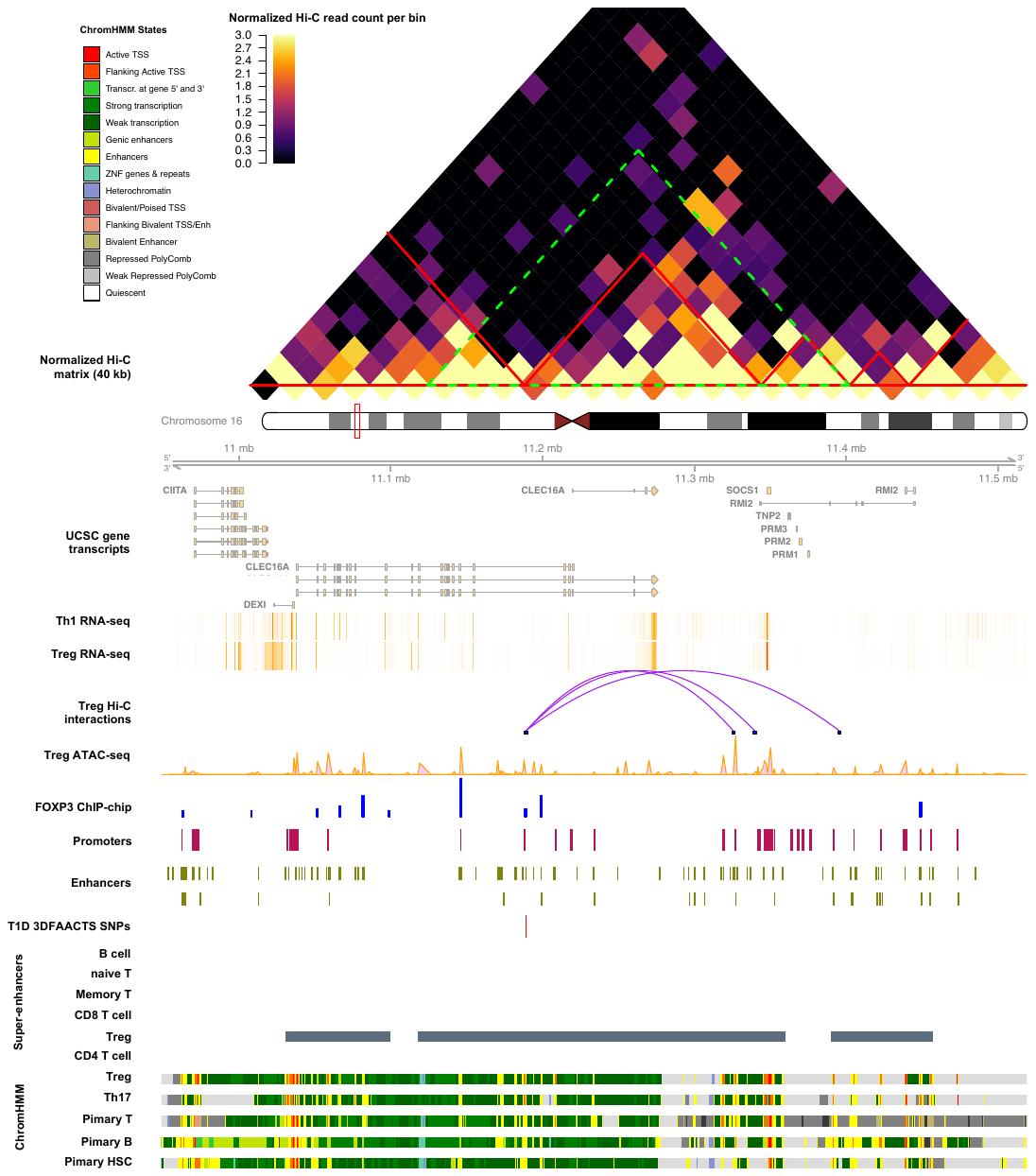

**Figure S10:** **Visualisation of the region of filtered T1D SNPs on chromosome 16.** Heatmap shows the Tregs Hi-C normalised interaction matrix (resolution of 40kb) on chr16: 10708949-11758949. The red triangles indicate Topologically Associated Domains (TADs) and the large green-dotted triangle indicates the boundary of the current plot. Tracks displayed below the chromosome 16 ideogram display workflow datasets (filtered SNPs, FOXP3-binding sites and Treg ATAC-seq, Hi-C interactions, promoters and enhancers) along with various types of cell type-specific data including UCSC Gene Transcript information, T cell subsets (Thelper1 and Treg) expression data, super-enhancer data and 15-state ChromHMM track of T cell lineages.

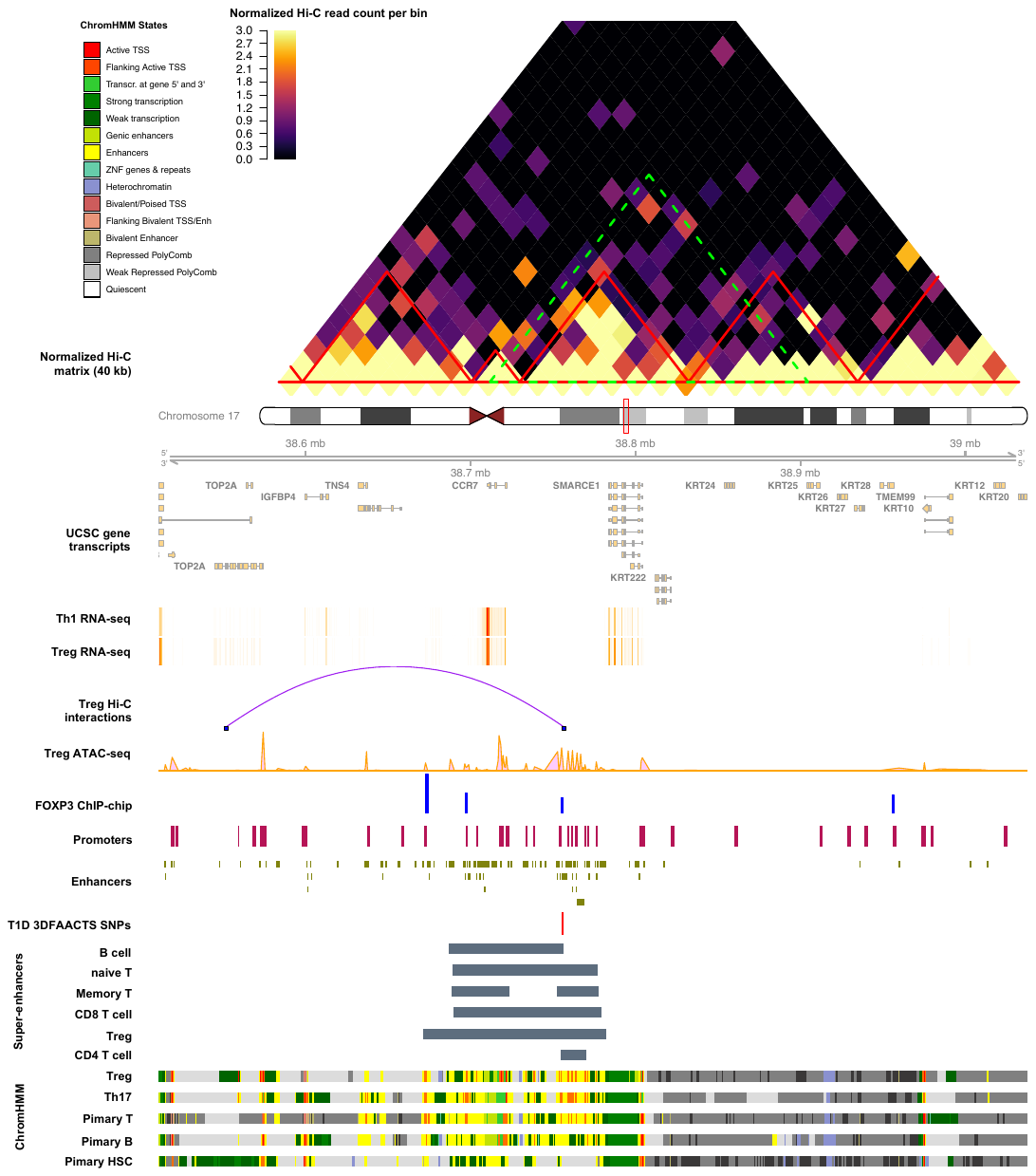

**Figure S11:** **Visualisation of the region of filtered T1D SNPs on chromosome 17.** Heatmap shows the Tregs Hi-C normalised interaction matrix (resolution of 40kb) on chr17: 38160811-39387558. The red triangles indicate Topologically Associated Domains (TADs) and the large green-dotted triangle indicates the boundary of the current plot. Tracks displayed below the chromosome 17 ideogram display workflow datasets (filtered SNPs, FOXP3-binding sites and Treg ATAC-seq, Hi-C interactions, promoters and enhancers) along with various types of cell type-specific data including UCSC Gene Transcript information, T cell subsets (Thelper1 and Treg) expression data, super-enhancer data and 15-state ChromHMM track of T cell lineages.

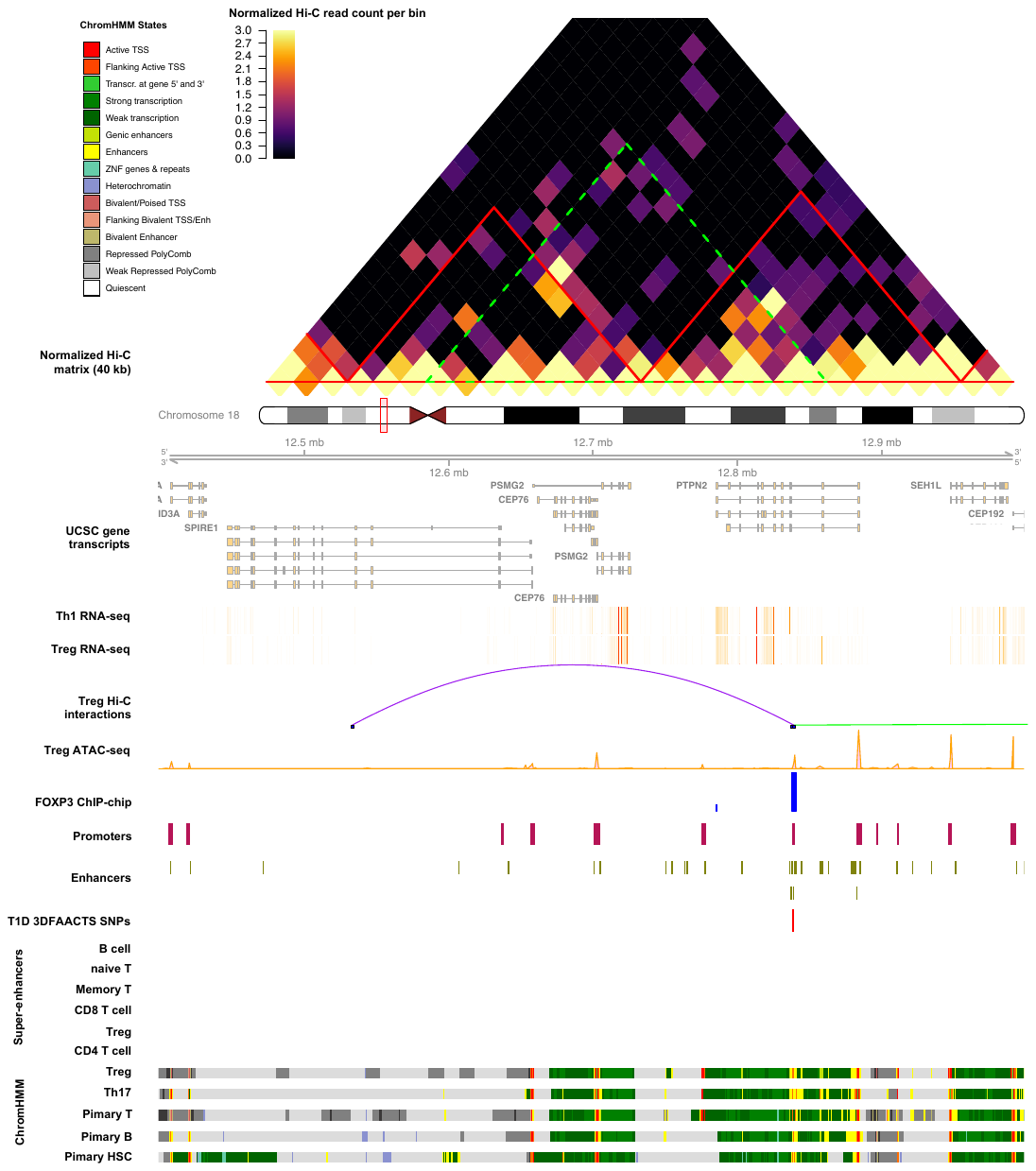

**Figure S12:** **Visualisation of the region of filtered T1D SNPs on chromosome 18.** Heatmap shows the Tregs Hi-C normalised interaction matrix (resolution of 40kb) on chr18: 12158767-13278767. The red triangles indicate Topologically Associated Domains (TADs) and the large green-dotted triangle indicates the boundary of the current plot. Tracks displayed below the chromosome 18 ideogram display workflow datasets (filtered SNPs, FOXP3-binding sites and Treg ATAC-seq, Hi-C interactions, promoters and enhancers) along with various types of cell type-specific data including UCSC Gene Transcript information, T cell subsets (Thelper1 and Treg) expression data, super-enhancer data and 15-state ChromHMM track of T cell lineages.

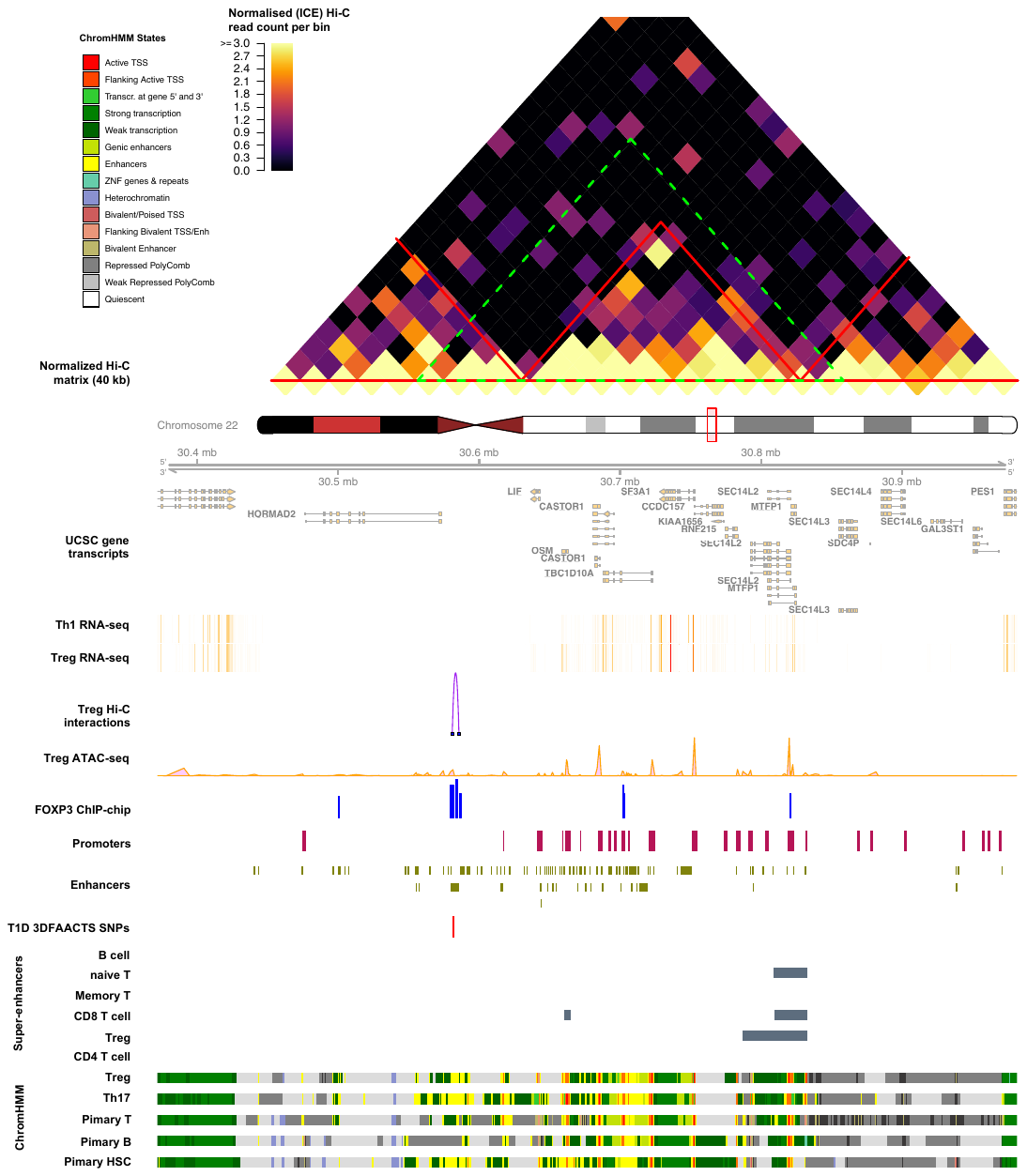

**Figure S13:** **Visualisation of the region of filtered T1D SNPs on chromosome 22.** Heatmap shows the Tregs Hi-C normalised interaction matrix (resolution of 40kb) on chr22: 30161722-31231722. The red triangles indicate Topologically Associated Domains (TADs) and the large green-dotted triangle indicates the boundary of the current plot. Tracks displayed below the chromosome 22 ideogram display workflow datasets (filtered SNPs, FOXP3-binding sites and Treg ATAC-seq, Hi-C interactions, promoters and enhancers) along with various types of cell type-specific data including UCSC Gene Transcript information, T cell subsets (Thelper1 and Treg) expression data, super-enhancer data and 15-state ChromHMM track of T cell lineages.

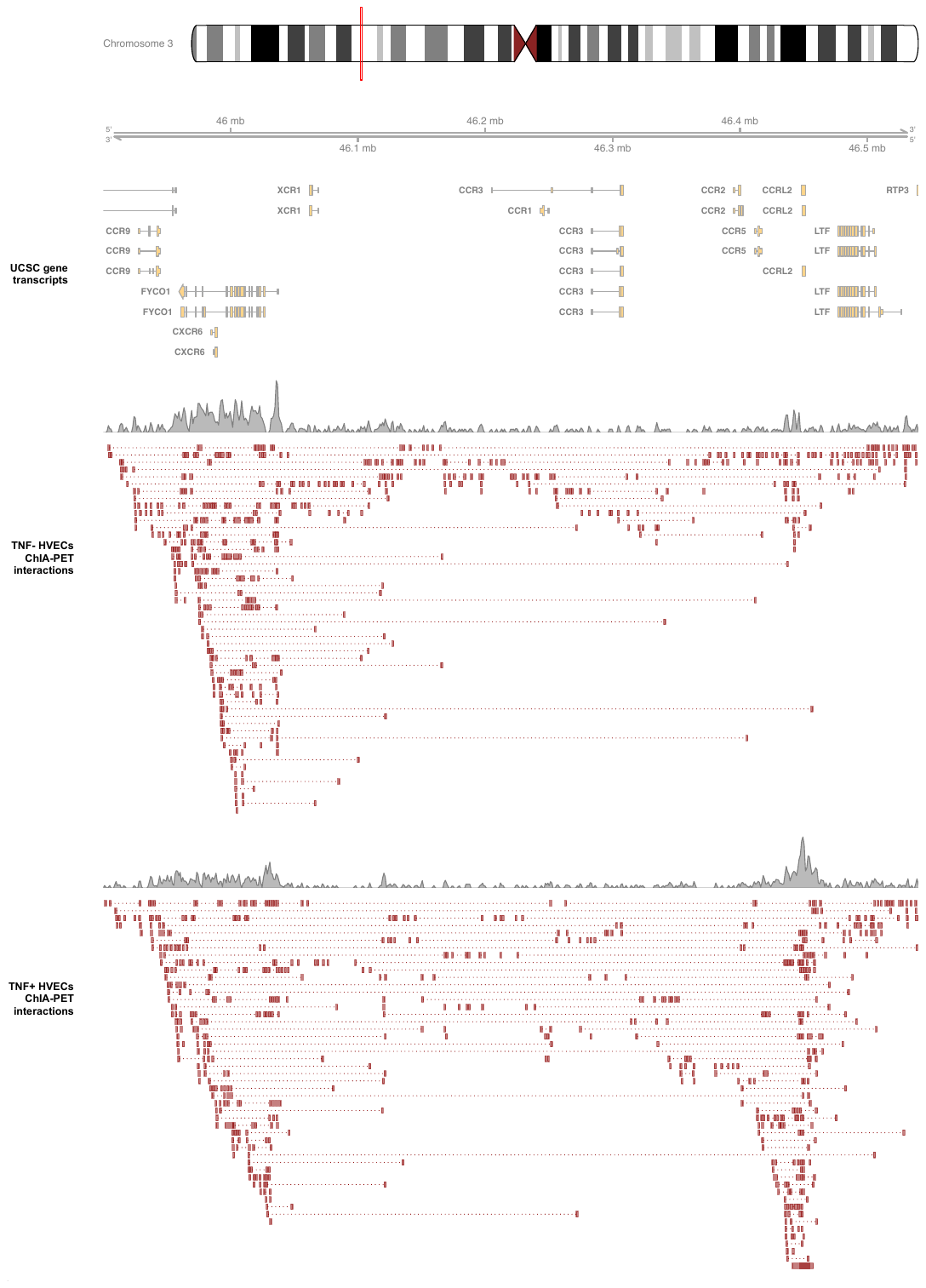

**Supplementary Figure 14:** **Published ChIA-PET data displaying the differences of interaction count after TNF induction in the CCR2/3/5 loci on chr3:45600000-46840000.**

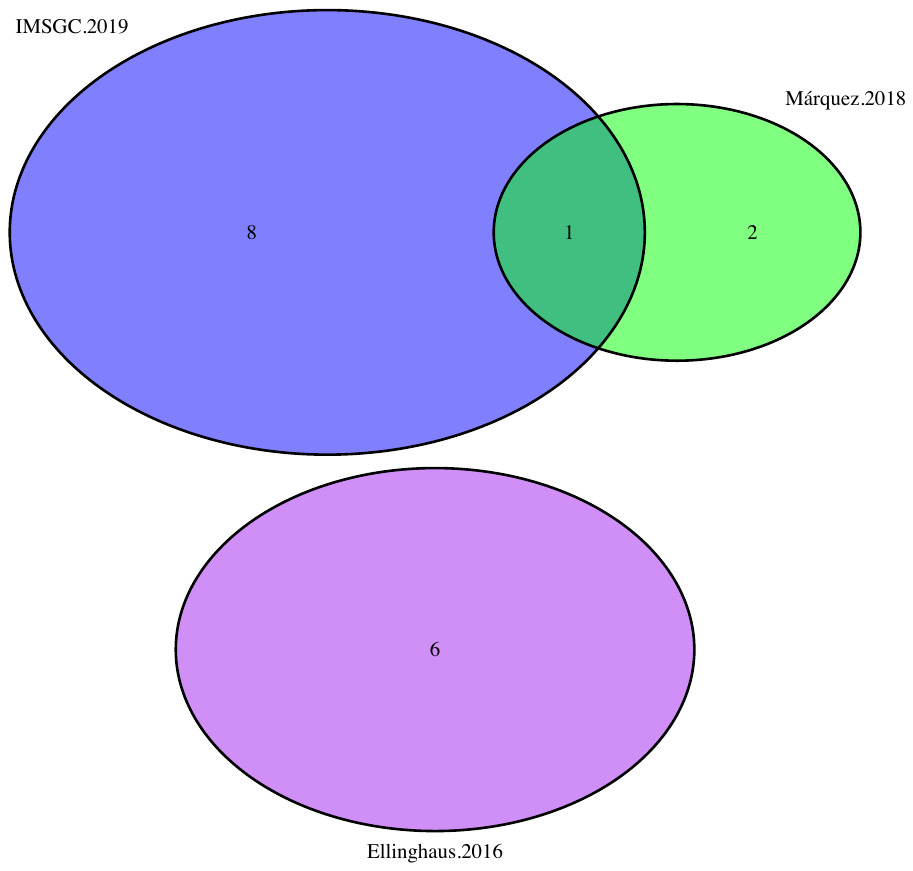

**Figure S15: Venn diagram showing shared filtered variants of 3 3DFAACTS SNP datasets from 3 different studies.** Three studies cover 10 autoimmune diseases, including T1D fine-mapped variants show in red, meta-analysed autoimmune variants of CeD, RA, SSc and T1D show in green, meta-analysed variants of five chronic inflammatory diseases, including AS, Crohn’s disease, psoriasis, PSC and UC show in purple, and fine-mapped MS variants show in blue.

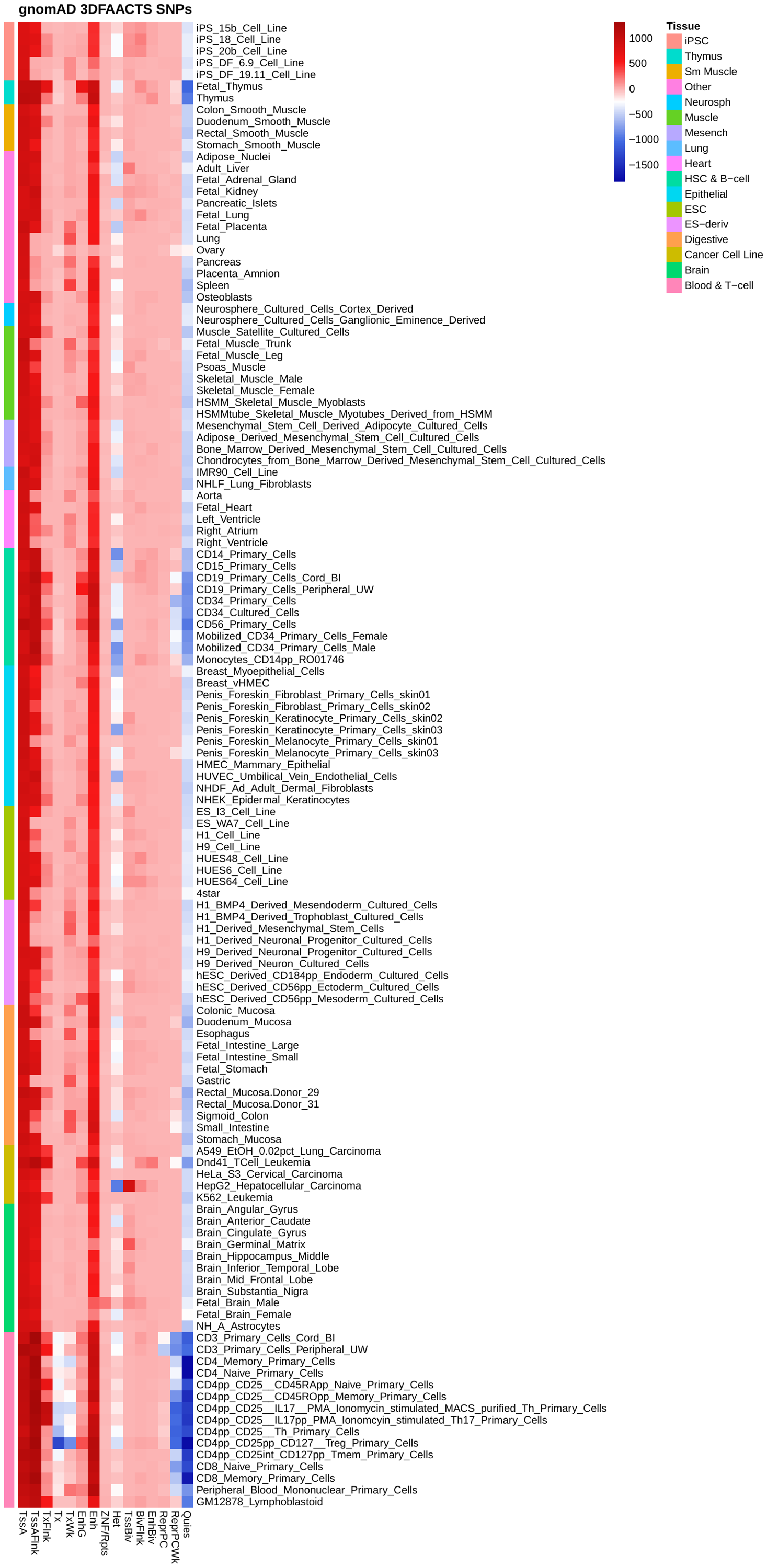

**Figure S16**: **Enrichment of filtered gnomAD variants found within NIH Epigenomics Roadmap samples.** Enrichment test of filtered gnomAD SNPs against chromHMM states from 129 tissues and cell types from Epigenomics Roadmap using GIGGLE. Red coloured regions indicate positive enrichment of variants within cell-types and chromHMM states, while blue coloured regions indicate negative enrichment.

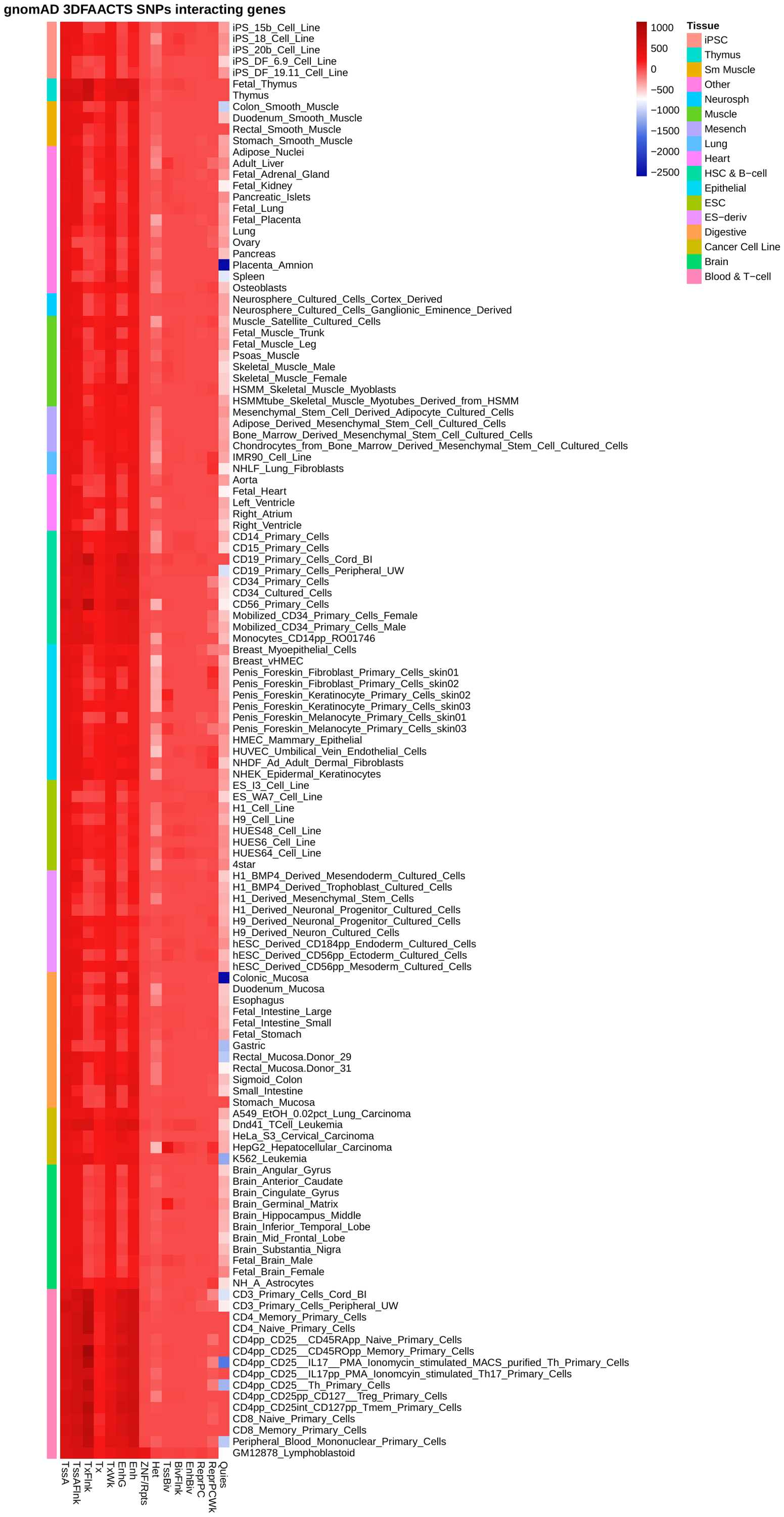

**Figure S17**: **Enrichment of 3D interacting regions of filtered gnomAD variants found within NIH Epigenomics Roadmap samples.** Enrichment test of filtered gnomAD SNPs against chromHMM states from 129 tissues and cell types from Epigenomics Roadmap using GIGGLE. Red coloured regions indicate positive enrichment of variants within cell-types and chromHMM states, while blue coloured regions indicate negative enrichment.

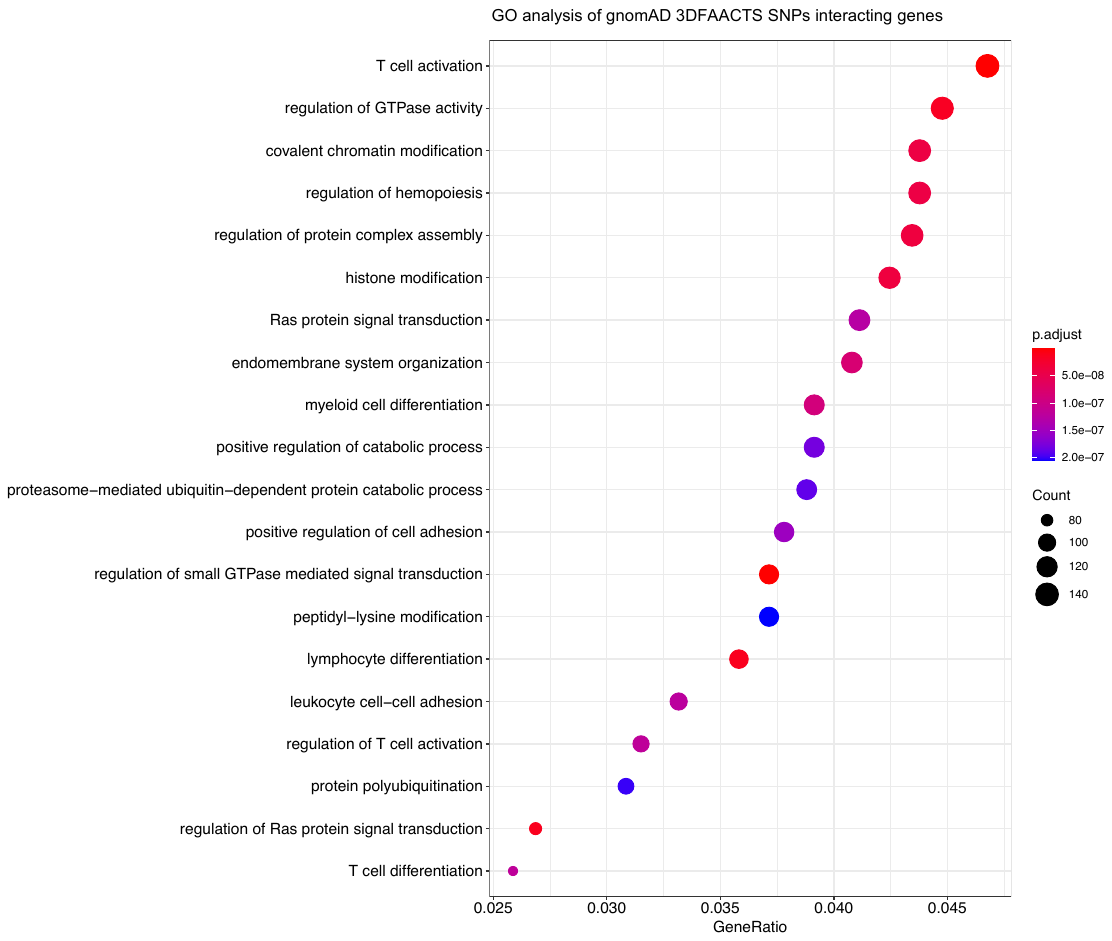

**Figure S18: Top20 significantly enriched gene ontology (GO) terms of common gnomAD 3DFAACTS SNPs 3D interacting genes.** Color indicates adjusted P-value of the enrichment test, and the sizes of the dots indicate the number of genes are included in the GO term.

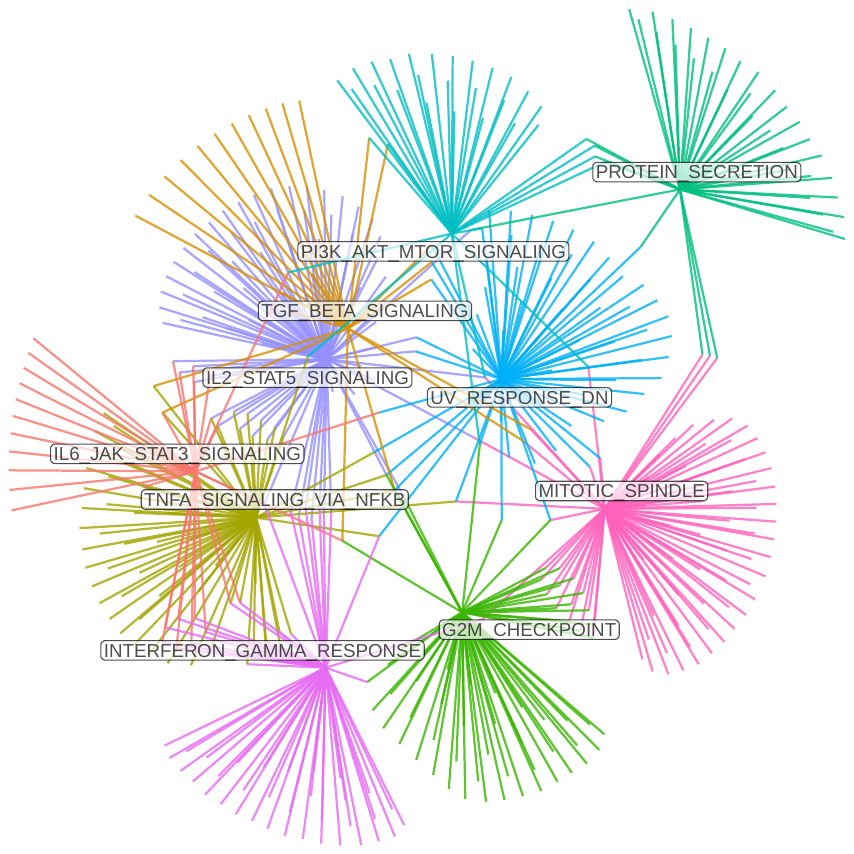

**Figure S19: GSEA network of nearest gene loci of filtered gnomAD SNPs using hallmark gene sets from MsigDB.** Larger colored nodes with name labels are enriched gene sets (FDR <= 0.05), smaller gray nodes are genes, edges indicate which genes are enriched in which gene sets.
